# Fluorescent Tools for Imaging and Ligand Screening of Dopamine D_2_-Like Receptors

**DOI:** 10.1101/2023.09.25.559398

**Authors:** Martin Nagl, Denise Mönnich, Niklas Rosier, Hannes Schihada, Alexei Sirbu, Nergis Konar, Irene Reyes-Resina, Gemma Navarro, Rafael Franco, Peter Kolb, Paolo Annibale, Steffen Pockes

**Affiliations:** Institute of Pharmacy, University of Regensburg, Universitätsstraße 31, D-93053, Regensburg, Germany; Department of Pharmaceutical Chemistry, University of Marburg, Marbacher Weg 6, 35037 Marburg, Germany; Max Delbrück Center for Molecular Medicine, Berlin 13125, Germany; CiberNed, Network Center for Neurodegenerative diseases, National Spanish Health Institute Carlos III, Madrid, Spain; Department Biochemistry and Physiology, School of Pharmacy and Food Sciences, Universitat de Barcelona, Barcelona, Spain; Department of Biochemistry and Molecular Biomedicine, Faculty of Biology, Universitat de Barcelona, Barcelona, Spain; School of Physics and Astronomy, University of St Andrews, North Haugh, St Andrews, Scotland; Department of Medicinal Chemistry, Institute for Therapeutics Discovery and Development, University of Minnesota, Minneapolis, MN 55414, USA

**Keywords:** Dopamine receptors, Fluorescent ligands, D_2_-like receptors, Confocal microscopy, Molecular brightness

## Abstract

The family of dopamine D_2_-like receptors represent an interesting target for a variety of neurological diseases, e.g. Parkinson’s disease (PD), addiction or schizophrenia. In this study we describe the synthesis of a new set of fluorescent ligands as tools for visualization of dopamine D_2_-like receptors. Pharmacological characterization in radioligand binding studies identified UR-MN212 (**20**) as a high-affinity ligand for D_2_-like receptors (pK_i_ (D_2long_R) = 8.24, pK_i_ (D_3_R) = 8.58, pK_i_ (D_4_R) = 7.78) with decent selectivity towards D_1_-like receptors. Compound **20** is a neutral antagonist in a G_o1_ activation assay at the D_2long_R, D_3_R and D_4_R, which is an important feature for studies using whole cells. The neutral antagonist **20**, equipped with a 5-TAMRA dye, displayed rapid association to the D_2long_R in binding studies using confocal microscopy demonstrating its suitability for fluorescence microscopy. Furthermore, in molecular brightness studies, the ligand’s binding affinity could be determined in a single-digit nanomolar range that was in good agreement with radioligand binding data. Therefore, the fluorescent compound can be used for quantitative characterization of native D_2_-like receptors in a broad variety of experimental setups.

## Introduction

Dopamine receptors are prominent members of the large family of G protein-coupled receptors (GPCRs) and are endogenously activated by the biogenic amine dopamine (Figure 1). Depending on their signaling pathways and sequence homology, dopamine receptors are divided into two families. The D_1_-like receptors (D_1_R, D_5_R) are mainly coupled to Gα_s_ and stimulate adenylyl cyclase (AC) upon activation while the D_2_-like receptors (D_2_R, D_3_R, D_4_R) mediate signal transduction predominantly via Gα_i/o_ which leads to inhibition of AC.^[1]^ Due to its high level of expression in the central nervous system (CNS) the D_2_-like family constitutes a highly interesting target for the treatment of several neurological diseases such as Parkinson’s disease (PD), addiction or schizophrenia.^[2,3]^ Especially the D_2_R is of utmost importance being the main target of agonists like pramipexole for the treatment of PD or antagonists like haloperidol for the therapy of schizophrenia (Figure 1).^[4,5]^ Despite intense research on both diseases in recent years the aforementioned drugs are still widely used in defiance of their approval dating back more than 25 years.^[6,7]^

**Figure 1.**
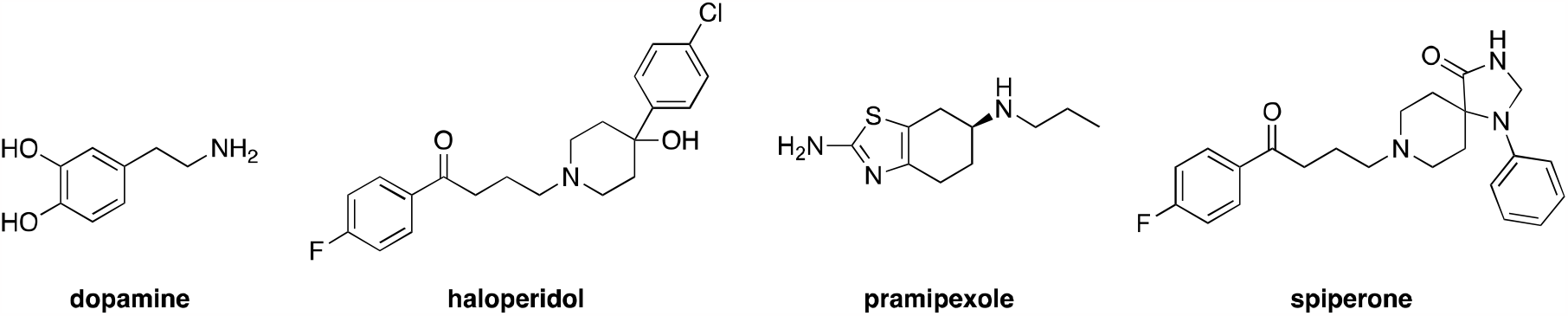
Chemical structures of dopamine, haloperidol, pramipexole and spiperone.

Evaluating binding affinities at the desired receptor is often one of the first steps concerning the development of new ligands. So far, binding properties have been mainly determined in radioligand competition binding experiments. Fluorescent ligands offer appealing alternatives to radioligands, especially, when it comes to targeting receptors using live cells and tissues or investigating the outcomes of drug-receptor interactions.^[8,9]^ A variety of fluorescence-based methods have been created to examine interactions between ligands and proteins, such as flow cytometry, fluorescence anisotropy, fluorescence polarization, fluorescence correlation spectroscopy and assays based on resonance energy transfer (RET).^[10–15]^ For these applications, fluorescence microscopy techniques such as confocal microscopy and total internal reflection fluorescence microscopy (TIRF) have emerged as state-of-the-art techniques to study receptor function.^[16–21]^

In this study, we report the synthesis of a small series of fluorescent ligands for D_2_-like receptors and their application in binding studies using molecular brightness in fluorescence microscopy. The aim of the study is to find valuable fluorescent tracers that can contribute to further exploration of the complex functions and interactions of the D_2_-like receptor family.

## Results and Discussion

### Design rationale

Fluorescent ligands share a common structure consisting of a pharmacophore, a linker region, and a fluorescent dye. Each of these components has a significant impact on the intended use of the fluorescent probe and must therefore be carefully selected. Spiperone was chosen as a pharmacophore (Figure 1) as it combines excellent affinity among D_2_-like receptors and high selectivity compared to D_1_ -like receptors.^[22–24]^ Its antagonistic mode of action makes it highly suitable for use in live cells since agonists might lead to receptor internalization.^[24]^ Moreover, additions of bulky structures to the aniline moiety have been described to be well tolerated in terms of affinity, making this part of the molecule a perfect attachment point for the linker.^[25,26]^ Linker length is a very important feature of a fluorescent ligand because it must allow the fluorescent dye to reach outside the binding pocket in order not to interfere with ligand binding. In contrast to previously reported N-(p-aminophenethyl)spiperone-based (NAPS) fluorescent ligands, where the dye was directly attached to the pharmacophore,^[25]^ we have designed two different linkers to gain more information about the necessary distance between pharmacophore and dye. A short linker based on γ-aminobutyric acid was designed to cover a rather short distance of 5 atoms to reduce the overall size of the ligand. The long linker covering a length of 18 atoms was based on polytheyleneglycole (PEG) units. This approach is very popular because PEG units are chemically stable, show only little interaction with cell membranes, and increase water solubility of the fluorescent probe.^[27]^ Since our goal was to design a fluorescent probe for microscopy we had to choose multilaterally usable dyes. Recent publications have reported that the 5-TAMRA dye is suitable for confocal microscopy and TIRF microscopy.^[13,28]^ As an alternative we selected DY-D549-P1 because it has similar spectral properties to 5-TAMRA. Since both dyes are hydrophilic, they are less prone to interact cell membranes than, for example, BODIPY fluorescent dyes, resulting in reduced nonspecific binding.^[29]^

### Chemistry

One of our aims was to find out how the selection of the respective dye and variation of linker length between the pharmacophore and the dye influence binding characteristics. Therefore, three different fluorescent ligands (**16, 17**, and **20**) differing in either the dye and/or linker length were designed. The synthesis of precursor NAPS (**11**) was carried out as previously described in ten steps (Scheme 1).^[26,30]^ In a first step intermediate **1** was synthesized from commercially available aniline and N-benzylpiperidin-4-one in the presence of HOAc and TMSCN. Subsequently the formed nitrile moiety was converted into an amide function using concentrated sulfuric acid leading to compound **2**. Reaction with DMF·DMA in methanol resulted in cyclisation to obtain 3. Reduction of the imidazolinone moiety with NaBH_4_ led to **4**. Cleavage of the benzyl group with ammoniumformiate in presence of Pd/C yielded **5**. Subsequently, an alkylation with 4-chloro-1-(4-fluorophenyl)butan-1-one in the presence of KI was performed to get **6**. At the same time, 2-(4-aminophenyl)ethan-1-ol (7) was Boc-protected to deliver **8** followed by a bromination with NBS to obtain **9**. Compound **10** was synthesized by N-alkylation of **6** with **9** in the presence of KOH and TBAB in toluene. TFA in DCM was used for deprotection of the aniline group to yield **11**.

In order to provide the fluorescent ligands with their dyes, corresponding spacers **12** and **13** were prepared for coupling with pharmacophore **11** by mono-Boc protection (Scheme 2A-B). For this purpose, 3,3’-((oxybis(ethane-2,1-diyl))bis(oxy))bis(propan-1-amine) and γ-aminobutyric acid were used as starting materials. For fluorescent ligand **16** and **17** succinic anhydride was added to the pharmacophoric scaffold **11** forming a terminal carbonic acid. This was subsequently coupled with spacer 12 (Scheme 2C) using HATU/DIPEA in DMF to yield intermediate **14**.^[25]^ Deprotection with TFA/DCM delivered precursor 15. The precursor for fluorescent ligand 20 was synthesized in a slightly different manner. Spacer 13 was directly coupled to 11 using HATU/DIPEA in DMF to obtain **18**, while cleavage of the Boc group under acidic conditions gave precursor **19**. In a final step, precursors **15** and **19** and the commercially available NHS-ester of 5-TAMRA or DY-D549-P1 were coupled in DMF in the presence of triethylamine.^[31]^ Purification with preparative HPLC afforded highly pure fluorescent ligands **16, 17**, and **20** (>98%) with great stability (Figure S1-S4; Supporting Information (SI)) in good to excellent yields (60-90 %).

**Scheme 1.**
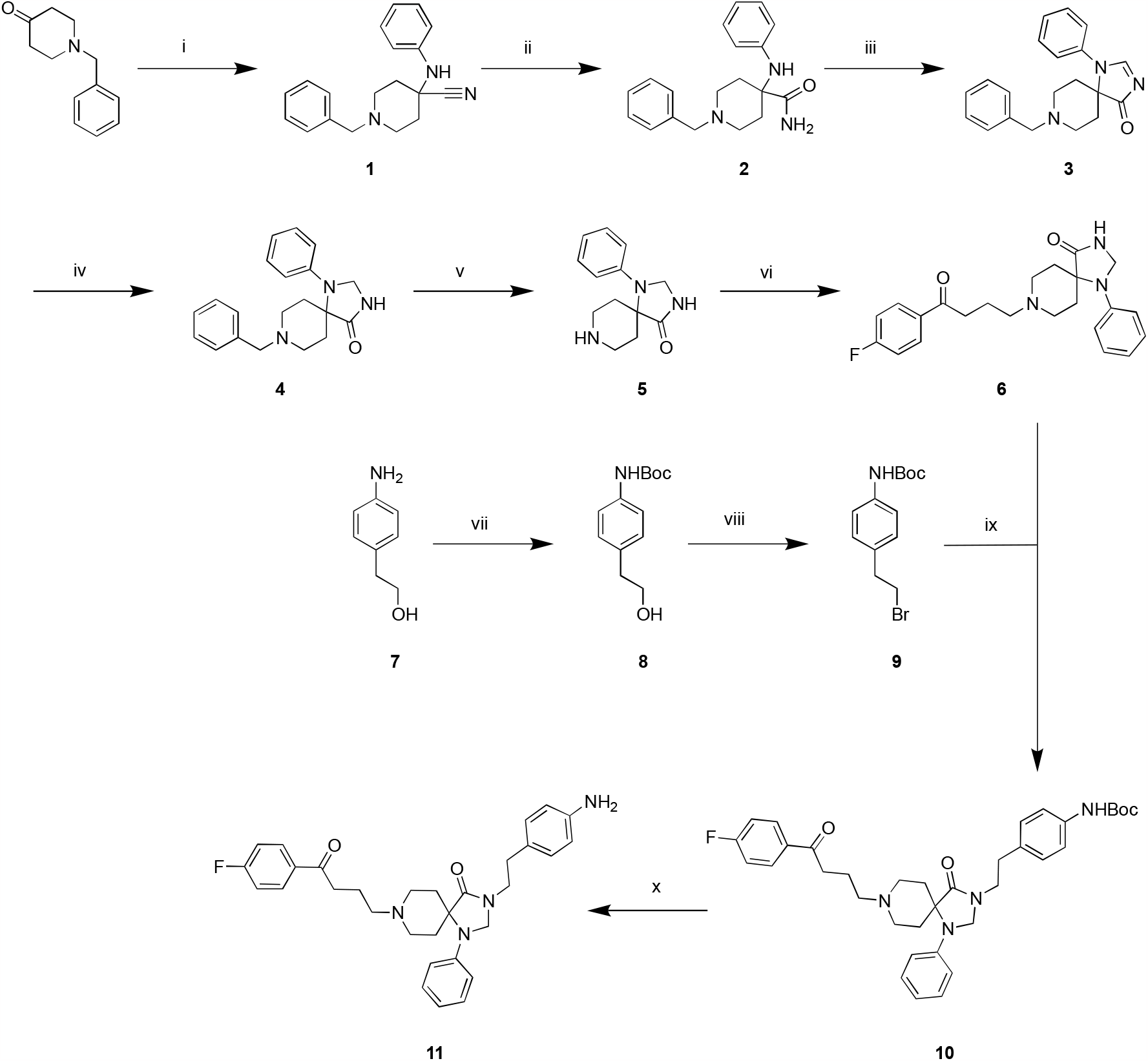
Synthesis of precursor N-(p-aminophenethyl)spiperone (NAPS) (**11**).^a^ ^a^Reactions and conditions: (i) aniline, TMSCN, HOAc, room temp, 4 h (99 %); (ii) H_2_SO_4_, room temp, 18 h (94 %); (iii) DMF-DMA, MeOH, 55°C, 16 h (75 %); (iv) NaBH_4_, MeOH, room temp, 4 h (70 %); (v) ammoniumformiate, Pd/C, MeOH, 55°C, 10 h (97 %); (vi) 4-chloro-1-(4-fluorophenyl)butan-1-one, Et_3_N, NaI, MeCN, reflux, 24 h (55 %); (vii) di-tert-butyl dicarbonate, AcOEt, room temp, 16 h (86 %); (viii) NBS, PPh_3_, DCM, 0°C, 3 h (88 %); (ix) KOH, TBAB, K_2_CO_3_, toluene, 90°C, 48 h (35 %); (x) TFA/DCM 1:4, room temp, overnight (83 %).

**Scheme 2.**
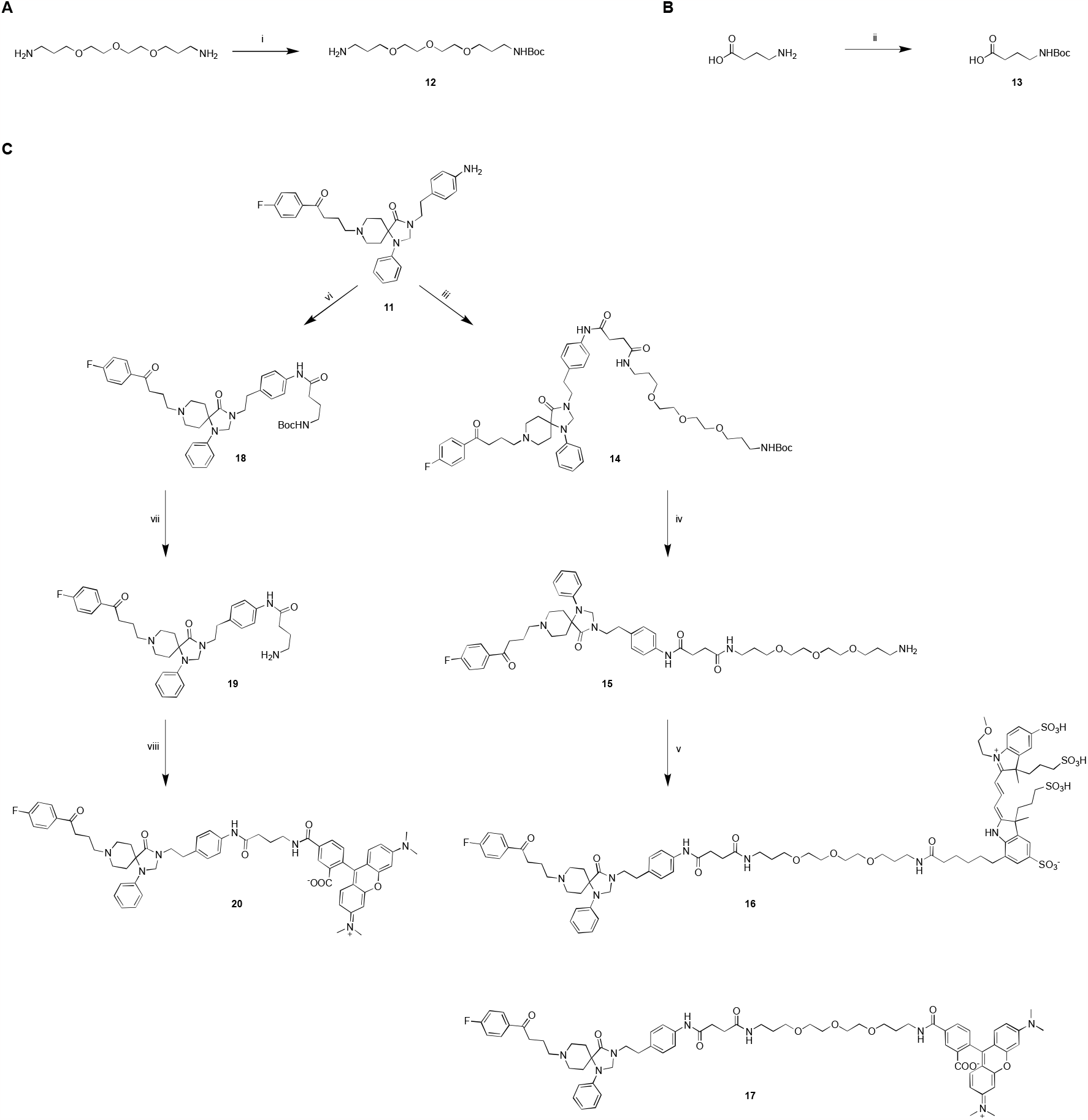
Synthesis of spacers **12** and **13** (**A. B**) and fluorescent ligands **16, 17**, and **20** (**C**).^a^ ^a^Reactions and conditions: (i) di-tert-butyl dicarbonate, Et_3_ N, DCM, rt, 5 h; (ii) di-tert-butyl dicarbonate, Et_3_N, DCM, rt, 10 h; (iii) a) succinic anhydride, DMF, rt, 10 h; b) **12**, HATU, DIPEA, DMF, rt, overnight; (iv) TFA/DCM 1:4, rt, 12 h; (v) 5-TAMRA NHS ester or DY-D549-P1 NHS ester, Et_3_N, DMF, rt, 4 h; (vi) **13**, HATU, DIPEA, DMF, overnight, rt; (vii) TFA/DCM 1:4, rt, 6 h.

### Pharmacological Characterization

In a first step, the synthesized compounds were tested for their binding properties to the D_2long_R. While 5-TAMRA labeled probes **17** and **20** showed very high affinity (p*K*_i_ = 8.25 and 8.24) for the target receptor in the single-digit nanomolar range, the addition of DY-D549-P1 to the pharmacophore resulted in a loss of affinity of 1.5 orders of magnitude (p*K*_i_ = 6.67). Subsequently, the selectivity of ligands **17** and **20** over the entire dopamine receptor family was determined. As expected, the selected compounds showed moderate to high affinity for the other D_2_R-like receptors, namely D_3_R (p*K*_i_ = 8.29 and 8.57) and D_4_R (p*K*_i_ = 7.53 and 7.78), whereas lower affinity for D_1_-like receptors was measured (cf. Figure 2; Table 1). In general, it could be observed that variations concerning the fluorescent dye had much more influence on binding affinities than different spacer lengths. Combined, compound **20** showed the highest affinities at all three D_2_-like receptors and we hence decided to use it as an representative of this fluorescent ligand series to determine its mode of action at D_2long_R, D_3_R, and D_4_R. We confirmed **20**’s antagonistic behavior using a BRET-based G_o1_ heterotrimer dissociation assay (G_o1_-CASE) (Figure 3, Table 2).^[32]^ A slight decrease was observed in the agonist mode only at high concentrations of 1 μM, most likely due to optical interference of **20** with the BRET components of G_o1_-CASE, as observed in a previous study with a 5-TAMRA-labeled histamine H_3_ receptor ligand (Figure 3A).^[13]^ In the antagonist-mode, however, **20** reduced the 1 μM dopamine-induced BRET response already at low nanomolar concentrations (p*K*_b_ (D_2long_R) = 10.53, p*K*_b_ (D_3_R) = 12.09, p*K*_b_ (D_4_R) = 8.78; Table 2; dopamine EC_50_ values required for the calculation of p*K*_b_ values were taken from the data shown in Figure S5, SI), further demonstrating that it acts as a neutral antagonist at D_2_-like receptors. This is of importance for microscopy studies, since agonistic properties of a ligand can lead to receptor internalization and subsequent degradation.

**Table 1.** Binding data of **16, 17** and **20** at the dopamine receptors.

| compound | Radioligand competition binding<sup>a</sup> |  |  |  |  |  |  |  |  |  |
| --- | --- | --- | --- | --- | --- | --- | --- | --- | --- | --- |
|  | p*K<sub>i</sub>* |  |  |  |  |  |  |  |  |  |
|  | D<sub>1</sub>R<sup>b</sup> | N | D<sub>2long</sub>R<sup>c</sup> | N | D<sub>3</sub>R<sup>d</sup> | N | D<sub>4</sub>R<sup>e</sup> | N | D<sub>5</sub>R<sup>f</sup> | N |
| **16** | n.d. | - | 6.67 ± 0.07 | 3 | n.d. | - | n.d. | - | n.d. | - |
| **17** | 6.48 ± 0.13 | 2 | 8.25 ± 0.03 | 3 | 8.29 ± 0.06 | 3 | 7.53 ± 0.04 | 3 | < 5.5 | 2 |
| **20** | 7.17 ± 0.09 | 2 | 8.24 ± 0.05 | 3 | 8.58 ± 0.16 | 3 | 7.78 ± 0.08 | 3 | 6.48 ± 0.04 | 2 |

<sup>a</sup>Competition binding assay at HEK293T\_CRE Luc 2P D<sub>1</sub>R homogenates, HEK293T\_CRE Luc 2P D<sub>2long</sub>R homogenates, HEK293T\_CRE Luc 2P D<sub>3</sub>R homogenates, HEK293T\_CRE Luc 2P D<sub>4</sub>R homogenates, or HEK293T\_CRE Luc 2P D<sub>5</sub>R homogenates. <sup>b</sup>Displacement of 1 nM [<sup>3</sup>H]SCH-23390 (*K<sub>d</sub>* = 0.3 nM). <sup>c</sup>Displacement of 0.05 nM [<sup>3</sup>H]*N*-Methylspiperone (*K<sub>d</sub>* = 0.015 nM). <sup>d</sup>Displacement of 0.05 nM [<sup>3</sup>H]*N*-Methylspiperone (*K<sub>d</sub>* = 0.026 nM). <sup>e</sup>Displacement of 0.1 nM [<sup>3</sup>H]*N*-Methylspiperone (*K<sub>d</sub>* = 0.078 nM). <sup>f</sup>Displacement of 1 nM [<sup>3</sup>H]SCH-23390 (*K<sub>d</sub>* = 0.4 nM). Data shown are mean values ± SEM of N experiments, each performed in triplicates. Data were analyzed by nonlinear regression and were best fitted to sigmoidal concentration-response curves. Displacement curves are shown in Figure 2.

**Table 2.** Functional data of **20** at the D_2_R, D_3_R and D_4_R.

| compound | Gα<sub>o1</sub> activation<sup>a</sup> |  |  |  |  |  |
| --- | --- | --- | --- | --- | --- | --- |
|  | p*K<sub>b</sub>* |  |  |  |  |  |
|  | D<sub>2long</sub>R<sup>b</sup> | N | D<sub>3</sub>R<sup>c</sup> | N | D<sub>4</sub>R<sup>d</sup> | N |
| **20** | 10.53 ± 0.06 | 3 | 12.09 ± 0.06 | 3 | 8.78 ± 0.17 | 3 |

<sup>a</sup>Competition binding experiment at HEK293 cells expressing the G<sub>o1</sub> BRET sensor with the wild type D<sub>2</sub>R, D<sub>3</sub>R or D<sub>4</sub>R. <sup>b</sup>Inhibition of dopamine-induced (c = 1 μM, EC<sub>50</sub> = 0.70 nM, Figure S5, SI) G<sub>o1</sub> activation. <sup>c</sup>Inhibition of dopamine-induced (c = 1 μM, EC<sub>50</sub> = 0.21 nM, Figure S5, SI) G<sub>o1</sub> activation. <sup>d</sup>Inhibition of dopamine-induced (c = 1 μM, EC<sub>50</sub> = 35 nM, Figure S5, SI) G<sub>o1</sub> activation. Data shown are mean values ± SEM of N experiments, each performed in duplicates. Data were analyzed by nonlinear regression and were best fitted to sigmoidal concentration-response curves. Displacement curves are shown in Figure 3.

**Figure 2:**
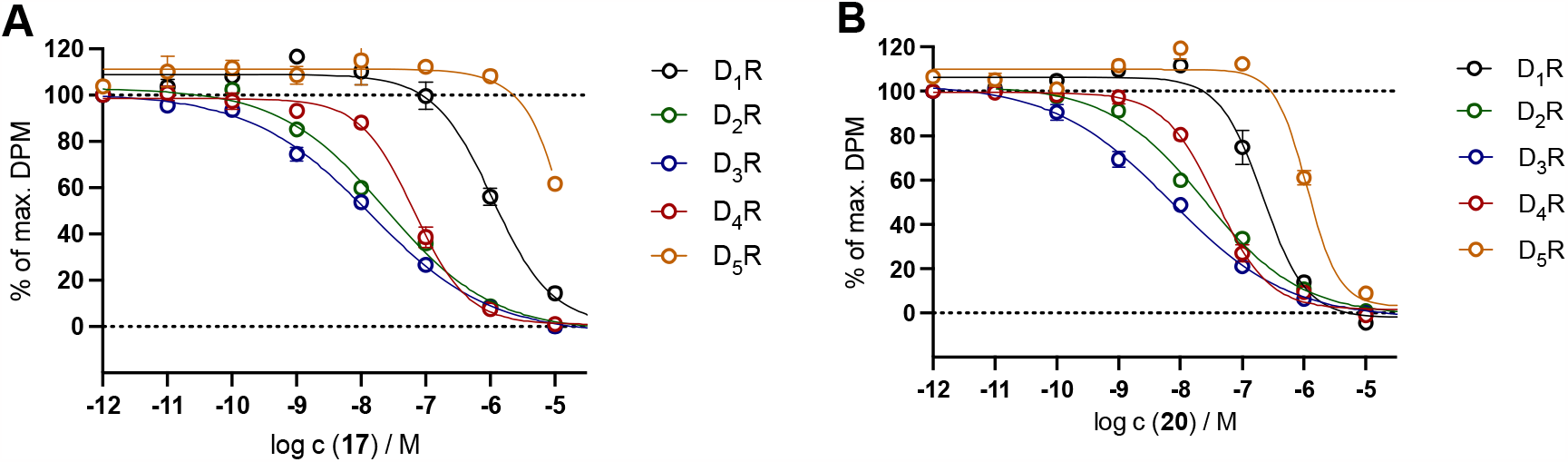
Displacement curves from radioligand competition binding experiments performed with **17** (**A**) or **20** (**B**) and the respective radioligands (cf. Table 1 footnotes) at the D_1-5_R. Graphs represent the means from N independent experiments each performed in triplicates. Data were analyzed by nonlinear regression and were best fitted to sigmoidal concentration-response curves.

**Figure 3.**
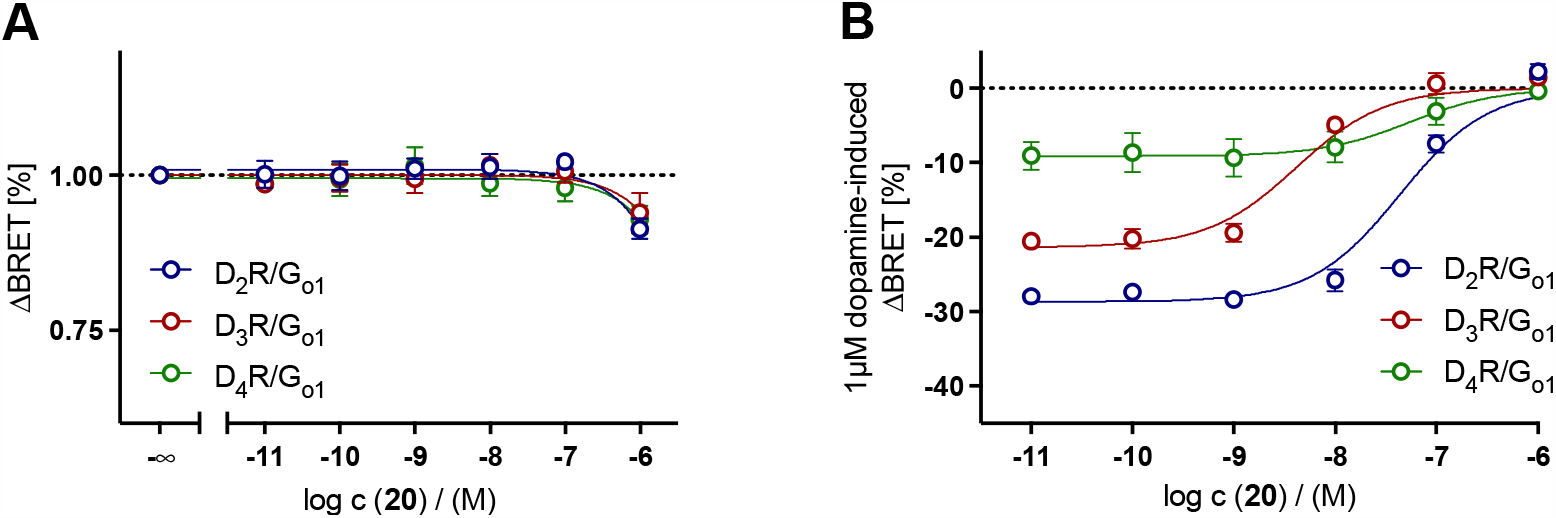
Concentration−response curves (CRCs) for G_o1_ activation of **20** in the absence (A) and presence (B) of 1 μM dopamine in HEK293A cells transiently expressing the G_o1_ BRET sensor along with the wild-type D_2_R, D_3_R or D_4_R. Graphs represent the means of three independent experiments each performed in duplicates. Data were analyzed by nonlinear regression and were best fitted to sigmoidal concentration-response curves.

### Fluorescence properties

Fluorescence excitation and emission spectra were recorded in order to further analyze the final compounds (in PBS containing 1% bovine serum albumin (BSA)) for their fluorescence properties. These are usually not greatly affected by the addition of a pharmacophore if the fluorophore is not chemically modified, as is the case with the dyes 5-TAMRA and DY549-P1. Nevertheless, the absorption and emission spectra as well as the quantum yield should be determined for the new fluorescent ligands. The emission and excitation spectra of 16, 17, and 20 are shown in Figure 4, while the excitation and emission maxima are presented in Table 3. The 5-TAMRA-labeled ligands 17 and 20 showed excitation maxima at 559/562 nm and emission maxima at 583/584 nm. Excitation maxima at 562 nm and emission maxima at 576 nm were recorded for the Dyomics-labeled compound 16. Quantum yields were determined in PBS + BSA 1 % with cresyl violet perchlorate as a red fluorescent standard according to a previously described procedure and are all in a good range of 36-39 % (Table 3).^[33]^ Measurements in PBS buffer, without 1 % BSA, gave decreased quantum yields, especially for compounds 16 and 17 (Table 3). Given these results, all ligands should be very well suitable for the use in fluorescence microscopy.

**Table 3.** Fluorescence properties of compounds **16, 17**, and **20**.

| compound | $\lambda_{\text{exc,max}}/\lambda_{\text{em,max}}$ (nm) | $\Phi$ (%)<sup>a</sup> | |
| --- | --- | --- | --- |
|  |  | PBS | PBS + 1 % BSA |
| **16** | 562 / 576 | 16.95 ± 0.52 | 37.48 ± 1.24 |
| **17** | 559 / 583 | 13.35 ± 0.38 | 38.47 ± 0.66 |
| **20** | 562 / 584 | 28.46 ± 0.41 | 36.27 ± 0.60 |

<sup>a</sup>Data shown are mean values ± SEM of three different slit adjustments (exc./em.): 5/5 nm, 5/10 nm, 10/10 nm.

**Figure 4.**
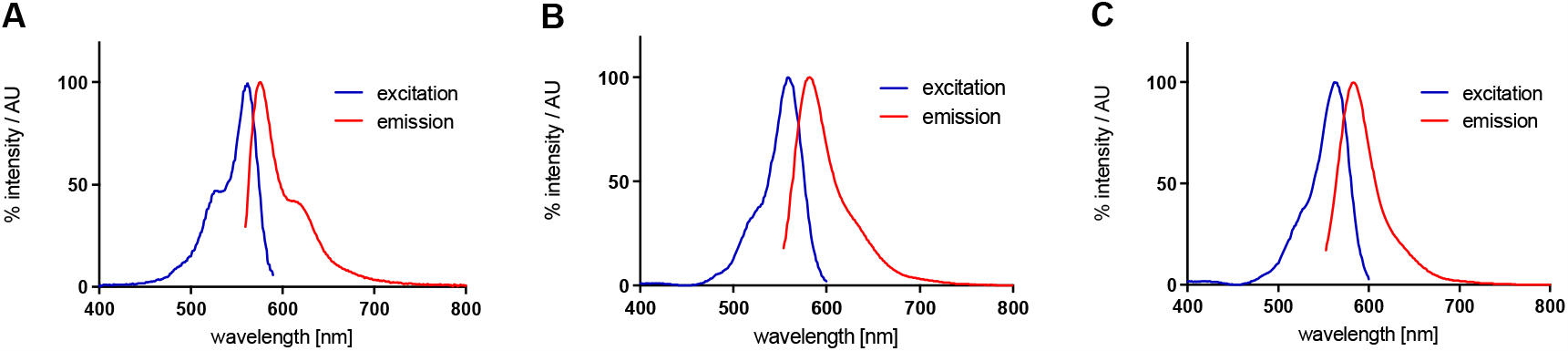
Excitation and emission spectra of **16** (**A**), **17** (**B**), and **20** (**C**) recorded in PBS buffer containing 1 % of BSA. The excitation wavelength for the emission spectra was 550 nm (**16**) or 545 nm (**17, 20**). The emission wavelength collected during the excitation scan was 590 nm (**16**) or 610 nm (**17, 20**).

### Microscopy

Confocal microscopy imaging was then used to visualize the binding behavior of compound **20**. Therefore, HEK293T cells were transiently transfected with DNA of the D_2long_R C-terminally fused to GFP_2_, allowing us to identify cells expressing high levels of the receptor (Figure 5A). Confocal microscopy images were acquired 48 hours after transfection (Figure 5). Once a suitable cell was identified, **20** (c = 50 nM) was added and rapid accumulation of fluorescence at the cell surface was observed (Figure 5B). This is due to the rapid association of **20** with the D_2long_R expressed on the cell membrane. These observations demonstrate the applicability of **20** for fluorescence microscopy experiments as a labeling tool to visualize D_2long_R in live cells.

**Figure 5:**
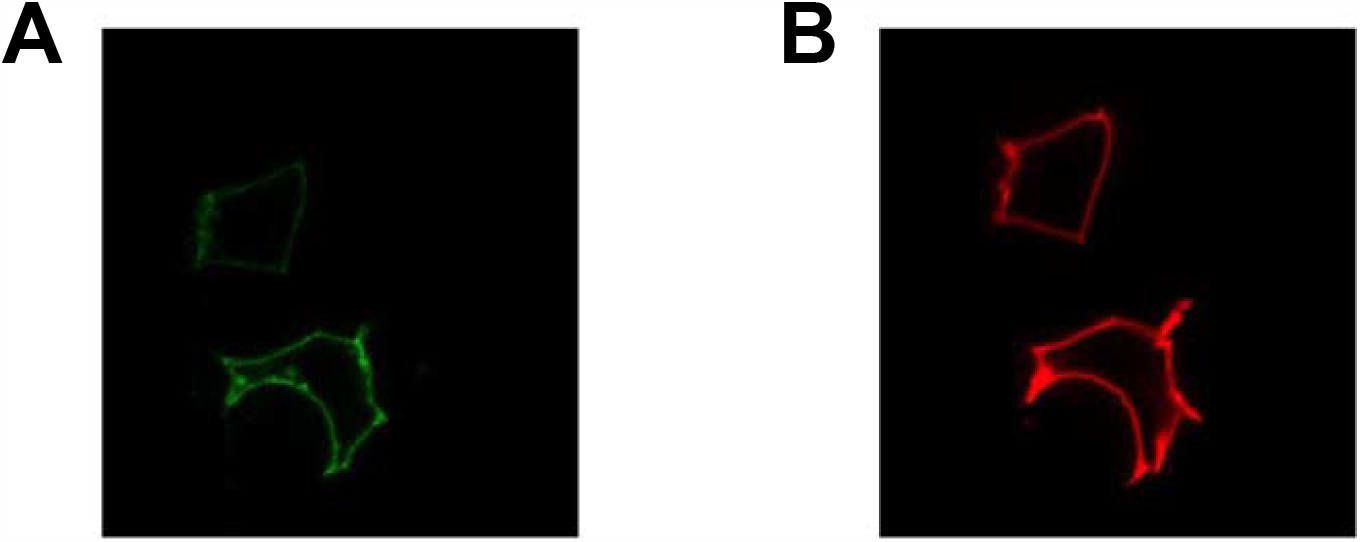
Confocal microscopy images: (**A**) Identification of cells expressing the D_2long_R-GFP_2_ receptor; (B) Fluorescence observed 3 min after addition of **20** (c = 50 nM).

### Molecular Brightness

We then used molecular brightness to quantify by an all-optical method the affinity of 20. When performed using two spectral channels, this approach allows correlating the number of receptors and fluorescent ligand at the membrane of intact, living cells.^[21]^ For this, HEK293-AD cells were transiently transfected with the C-terminally tagged D_2long_R-mNeonGreen and incubated with different concentrations of 5-TAMRA-labeled **20**. After allowing equilibration, the ligand was washed out to avoid background of freely diffusing ligand and basolateral membranes were immediately imaged on a confocal microscope (Figure 6A). The photon counts per pixel and their variance in a region of interest were used to calculate the average number of fluorescent emitters per confocal excitation volume and their brightness. The emitter numbers were calculated for both mNeonGreen (receptor) and for 5-TAMRA (ligand) channels and plotted against each other. Figure 6B shows the values and superposed a linear regression fit (Figure 6B). A slope of 0.80 ± 0.03 obtained for 575 nM of **20** indicates a high, albeit partial, occupancy of the receptor by the ligand. Likely reasons for not observing full receptor occupancy could be receptor signal from areas within the confocal volume and proximal to the membrane (e.g. early endosomes), quick ligand dissociation kinetics or slight photobleaching of the TAMRA. Nonetheless, the concentration response curve, formed by the slopes plotted against the corresponding ligand concentrations, yields a pK_d_ value of 8.12 ± 0.11 mirroring the radioligand binding data (Figure 6C).

**Figure 6.**
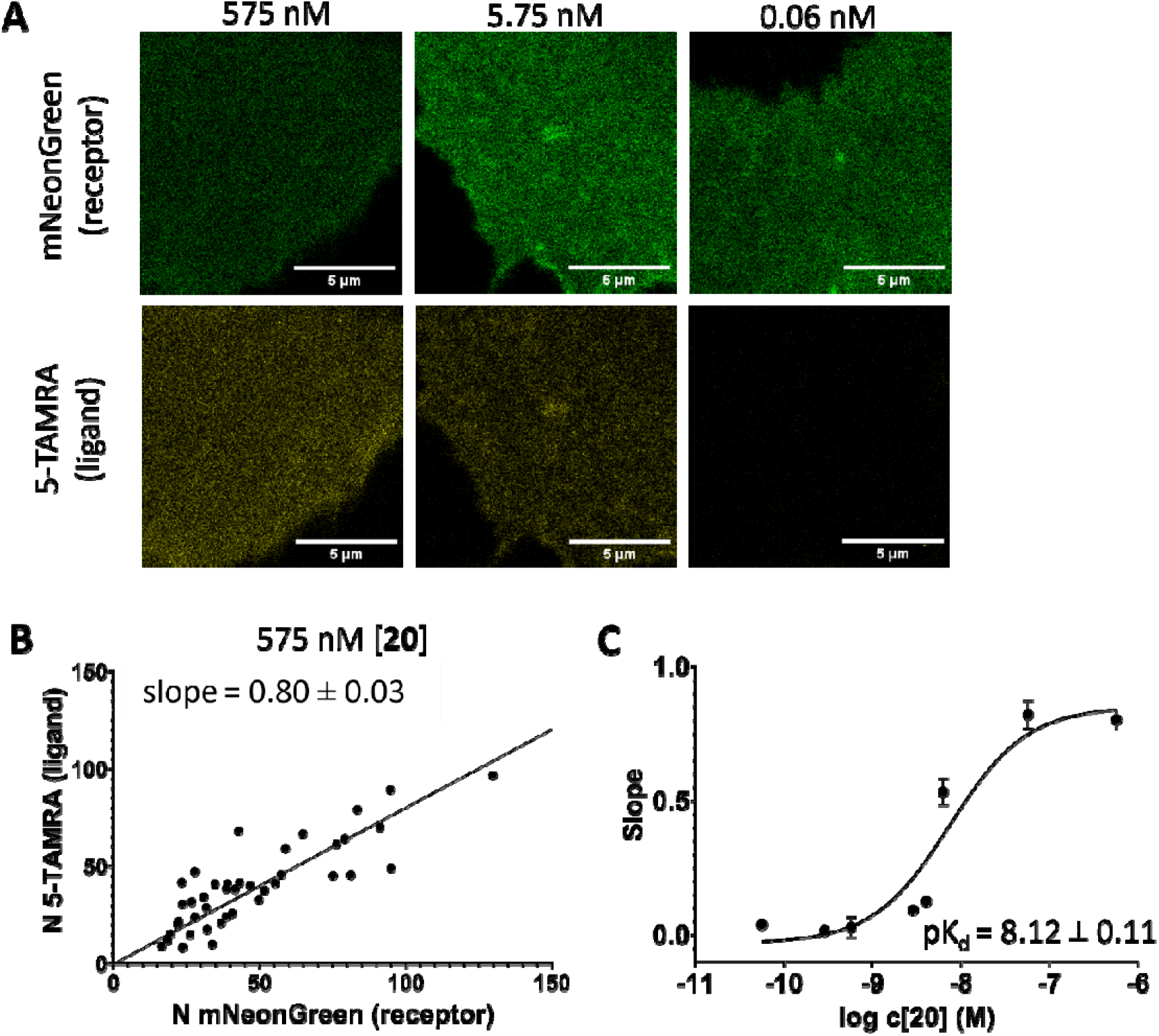
Association of **20** to the hD_2_R using molecular brightness analysis. Basolateral membranes of HEK-293AD cells transiently expressing hD_2_R-mNeonGreen and preincubated with indicated amount of **20** (**A**); calculated numbers of emitters for 5-TAMRA and mNeonGreen channels, and the corresponding linear regression fit (mean ± SEM; n = 41 cells from 5 independent experiments) (**B**), slopes obtained from linear fits plotted against log concentrations of **20** with the corresponding non-linear fit and pK_d_ value (mean ± SEM; n = at least 18 cells from 3 independent experiments for each datapoint) (**C**).

### Conclusion

In this study, a series of three fluorescent ligands were designed and synthesized containing different linkers and fluorescent dyes. The well-known antagonist spiperone was selected as a scaffold as it combines excellent affinity among D_2_-like receptors and selectivity towards D_1_R and D_5_R, while 5-TAMRA and DY549-P1 were chosen as fluorescent dyes proposing suitability as fluorescent tracer for microscopy studies. In a 13- and 14-step synthesis, respectively, the fluorescent ligands were successfully prepared and purified for further characterization. Radioligand binding studies revealed that both 5-TAMRA ligands, **17** (p*K*_i_ (D_2long_R) = 8.25, p*K*_i_ (D_3_R) = 8.29, p*K*_i_ (D_4_R) = 7.53) and **20** (p*K*_i_ (D_2long_R) = 8.24, p*K*_i_ (D_3_R) = 8.58, p*K*_i_ (D_4_R) = 7.78), have high-affinities to D_2_-like receptors with good fluorescence properties and quantum yield. In a BRET-based G_o1_ heterotrimer dissociation assay, the expected antagonistic mode of action of fluorescent ligand **20** at the D_2long_R, D_3_R and D_4_R could be determined, demonstrating the suitability of **20** for whole cell applications, e.g. fluorescence microscopy. In addition to successful association experiments in confocal microscopy, molecular brightness studies confirmed ligand binding to the D_2_R in the single-digit nanomolar range. In conclusion, our study provides an interesting set of new fluorescent ligands for D_2_-like receptors, which can be applied in various applications in fluorescence microscopy in the future.

### Experimental Section

#### Chemistry

Commercially available chemicals and solvents were purchased from standard commercial suppliers (Merck (Darmstadt, Germany), Sigma-Aldrich (Munich, Germany), Acros Organics (Geel, Belgium), Alfa Aesar (Karlsruhe, Germany), abcr (Karlsruhe, Germany) or TCI Europe (Zwijndrecht, Belgium) and were used as received. All solvents were of analytical grade. The fluorescent dye 5-TAMRA NHS ester was purchased from Lumiprobe (Hannover, Germany), the fluorescent dye DY-549 P1 NHS ester was purchased form Dyomics GmbH (Jena, Germany). Deuterated solvents for nuclear magnetic resonance (^1^H NMR and ^13^C NMR) spectra were purchased from Deutero GmbH (Kastellaun, Germany). All reactions carried out with dry solvents were accomplished in dry flasks under nitrogen or argon atmosphere. For the preparation of buffers, HPLC eluents, and stock solutions millipore water was used. Column chromatography was accomplished using Merck silica gel Geduran 60 (0.063-0.200 mm) or Merck silica gel 60 (0.040-0.063 mm) (flash column chromatography). The reactions were monitored by thin layer chromatography (TLC) on Merck silica gel 60 F254 aluminium sheets and spots were visualized under UV light at 254 nm, by potassium permanganate or ninhydrin staining. Lyophilization was done with a Christ alpha 2-4 LD equipped with a vacuubrand RZ 6 rotary vane vacuum pump. Nuclear magnetic resonance (^1^H NMR and ^13^C NMR) spectra were recorded on a Bruker (Karlsruhe, Germany) Avance 300 (^1^H: 300 MHz, ^13^C: 75 MHz) or 400 (^1^H: 400 MHz, ^13^C: 101 MHz) spectrometer using perdeuterated solvents. The chemical shift δ is given in parts per million (ppm). Multiplicities were specified with the following abbreviations: s (singlet), d (doublet), t (triplet), q (quartet), quin (quintet), m (multiplet), and br (broad signal) as well as combinations thereof. ^13^C NMR-Peaks were determined by DEPT 135 and DEPT 90 (distortionless enhancement by polarization transfer). NMR spectra were processed with MestReNova 11.0 (Mestrelab Research, Compostela, Spain). High-resolution mass spectrometry (HRMS) was performed on an Agilent 6540 UHD Accurate-Mass Q-TOF LC/MS system (Agilent Technologies, Santa Clara, CA) using an ESI source. Preparative HPLC was performed with a system from Waters (Milford, Massachusetts, USA) consisting of a 2524 binary gradient module, a 2489 detector, a prep inject injector and a fraction collector III. A Phenomonex Gemini 5μm NX-C18 column (110 Å, 250 x 21.2 mm, Phenomenex Ltd., Aschaffenburg, Germany) served as stationary phase. As mobile phase, 0.1% TFA (method A) or 0.1% NH_3_ (method B) in millipore water and acetonitrile (MeCN) were used. The temperature was 25 °C, the flow rate 20 mL/min and UV detection was performed at 560 nm. Analytical HPLC experiments were performed on a 1100 HPLC system from Agilent Technologies equipped with Instant Pilot controller, a G1312A Bin Pump, a G1329A ALS autosampler, a G1379A vacuum degasser, a G1316A column compartment and a G1315B DAD detector. The column was a Phenomenex Gemini 5μm NX-C18 column (110 Å, 250 x 4.6 mm, Phenomenex Ltd., Aschaffenburg, Germany), tempered at 30 °C. As mobile phase, mixtures of MeCN and aqueous TFA (method A) or aqueous NH_3_ (method B) were used (linear gradient: MeCN/TFA or NH_3_ (0.1%) (v/v) 0 min: 10:90, 25-35 min: 95:5, 36-45 min: 10:90; flow rate = 1.00 mL/min, t_0_ = 3.21 min). Capacity factors were calculated according to k = (t_R_ -t_0_)/t_0_. Detection was performed at 220 and 254 nm. Furthermore, a filtration of the stock solutions with PTFE filters (25 mm, 0.2 μm, Phenomenex Ltd., Aschaffenburg, Germany) was carried out before testing. Compound purities determined by HPLC were calculated as the peak area of the analyzed compound in % relative to the total peak area (UV detection at 220 or 254 nm). The HPLC purity and stability of final compounds is displayed in the SI (Figure S1-S4, SI).

#### Synthesis and Analytical Data

##### 1-Benzyl-4-(phenylamino)piperidine-4-carbonitrile (1)^[30]^

TMSCN (2.97 g, 30.0 mmol, 3 eq) was added dropwise to a solution of aniline (0.93 g, 10.0 mmol, 1 eq) and N-benzyl-piperidone (1.89 g, 10.0 mmol, 1 eq) in glacial acid (15 mL) at 0°C. The reaction was stirred at room temperature for 5 h. After the reaction was complete, monitored by TLC, the solution was basified with aqueous KOH 20% and extracted with DCM (3 x 30 mL). The combined organic phases were dried over Na_2_SO4 and the solvent was removed under reduced pressure. Column chromatography (DCM/MeOH 99/1) afforded **2** (3.01 g, 98%) as a yellow foam. ^1^H NMR (400 MHz, CDCl_3_) δ 7.32 – 7.24 (m, 4H), 7.25 – 7.15 (m, 3H), 6.89 – 6.82 (m, 3H), 3.62 (s, 1H), 3.50 (s, 2H), 2.82 – 2.67 (m, 2H), 2.47 – 2.36 (m, 2H), 2.32 – 2.21 (m, 2H), 1.93 – 1.81 (m, 2H). ^13^C NMR (101 MHz, CDCl_3_) δ 143.37, 137.92, 129.33, 129.06, 128.95, 128.40, 127.33, 120.95, 120.74, 117.82, 62.61, 53.10, 49.32, 36.10. HRMS (ESI-MS): m/z [M+H^+^] calculated for C_19_ H_22_ N_3_^+^: 292.1808, found 292.1813; C_19_H_21_N_3_ (291.40).

##### 1-Benzyl-4-(phenylamino)piperidine-4-carboxamide (2)^[30]^

**2** (3.00 g, 10.0 mmol) was dissolved in 10 mL H_2_SO_4_ (96% w/w) and stirred at room temperature overnight. The reaction was then carefully basified (pH>10) by dropwise addition of aqueous NaOH (30%) while maintaining the temperature below 0°C. The resulting mixture was extracted with DCM (3 x 30 mL). The organic phases were combined and dried over Na_2_SO_4_. The solvent was removed under reduced pressure to yield **3** (2.99 g, 94%) as an orange oil. ^1^H NMR (300 MHz, DMSO-d_6_) δ 7.34 – 7.17 (m, 6H), 7.11 – 7.00 (m, 3H), 6.67 – 6.53 (m, 3H), 5.49 (s, 1H), 3.43 (s, 2H), 2.59 – 2.48 (m, 2H), 2.26 (t, J = 10.5 Hz, 2H), 2.10 – 1.95 (m, 2H), 1.93 – 1.77 (m, 2H). ^13^C NMR (101 MHz, DMSO-d_6_) δ 178.14, 145.93, 139.16, 129.16, 128.94, 128.61, 127.28, 116.94, 115.17, 62.70, 57.53, 48.94, 31.84. HRMS (ESI-MS): m/z [M+H^+^] calculated for C_19_ H_24_ N_3_ O^+^: 310.1914, found 310,19120; C_19_ H_23_ N_3_ O (309.41).

##### 8-Benzyl-1-phenyl-1,3,8-triazaspiro[4.5]dec-2-en-4-one (3)^[30]^

DMF-DMA (3.44 g, 29.1 mmol, 3 eq) was added to a solution of **3** (2.99 g, 9,7 mmol, 1 eq) in methanol (40 mL) and the mixture was stirred for 16 h at 55°C. The solvent was evaporated and the crude residue was purified by column chromatography (DCM/MeOH 95/5) to yield **4** (3.01 g, 96%) as a yellow semi solid.^1^H NMR (300 MHz, CDCl_3_) δ 8.22 (s, 1H), 7.53 – 7.43 (m, 3H), 7.32 – 7.20 (m, 5H), 7.19 – 7.14 (m, 2H), 3.56 (s, 2H), 3.08 – 2.94 (m, 2H), 2.72 – 2.59 (m, 2H), 2.05 – 1.93 (m, 2H), 1.84 – 1.71 (m, 2H). ^13^C NMR (75 MHz, CDCl_3_) δ 194.06, 169.16, 135.38, 130.01, 129.58, 129.16, 128.26, 128.06, 127.11, 65.05, 62.62, 46.83, 30.88. HRMS (ESI-MS): m/z [M+H^+^] calculated for C_20_H_22_N_3_O^+^: 320.1757, found 320.1765; C_20_H_21_N_3_O (319.41).

##### 8-Benzyl-1-phenyl-1,3,8-triazaspiro[4.5]decan-4-one (4)^[30]^

NaBH_4_ (0.45 g, 11.8 mmol, 1.25 eq) was added to a solution of **4** (3.00 g, 9.4 mmol, 1 eq) in methanol (35 mL). The reaction was stirred at room temperature whereupon a white solid precipitated. After the reaction was complete, monitored by TLC, the solid was filtered off to obtain **5** (2.08 g, 70%) as a white foam. ^1^H NMR (300 MHz, DMSO-d_6_) δ 8.63 (s, 1H), 7.38 – 7.31 (m, 4H), 7.29 – 7.21 (m, 3H), 6.87 (d, J = 8.1 Hz, 2H), 6.76 (t, J = 7.3 Hz, 1H), 4.57 (s, 2H), 3.52 (s, 2H), 2.76 – 2.67 (m, 4H), 2.59 – 2.52 (m, 2H), 1.57 (d, J = 13.5 Hz, 2H). ^13^C NMR (75 MHz, DMSO-d_6_) δ 176.51, 143.78, 139.18, 129.53, 129.17, 128.67, 127.30, 118.16, 114.72, 62.62, 58.64, 58.61, 49.71, 28.88. HRMS (ESI-MS): m/z [M+H^+^] calculated for C_20_ H_24_ N_3_ O^+^: 322.1914, found 322.1920; C_20_ H_23_ N_3_ O (321.42).

##### 1-Phenyl-1,3,8-triazaspiro[4.5]decan-4-one (5)^[30]^

**5** (2.08 g, 6.5 mmol, 1 eq) was dissolved in methanol (50 mL) and glacial acid (1 mL). To the obtained solution a catalytic amount of palladium on activated charcoal (10 % Pd basis) and ammoniumformiate (1.60 g, 26.0 mmol, 4 eq) was added. The mixture was then stirred at 55°C until the reaction was complete, indicated by TLC. The mixture was filtered through Celite, concentrated under reduced pressure, diluted with water and basified with aqueous KOH (20%). The aqueous phase was extracted with EtOAc (3 x 30 mL). The combined organic layers were dried over Na_2_SO_4_ and the solvent was removed under reduced pressure. **6** (1.45 g, 97%) was obtained as a colorless oil. ^1^H NMR (400 MHz, DMSO-d_6_) δ 8.58 (s, 1H), 7.26 – 7.15 (m, 2H), 6.88 (d, J = 8.3 Hz, 2H), 6.71 (t, J = 7.3 Hz, 1H), 4.56 (s, 2H), 3.18 – 3.07 (m, 4H), 2.83 (dd, J = 12.2, 4.9 Hz, 2H), 2.39 (td, J = 13.0, 5.5 Hz, 2H), 1.46 (d, J = 13.4 Hz, 2H). ^13^C NMR (101 MHz, DMSO-d_6_) δ 176.92, 143.89, 129.39, 117.69, 114.39, 59.24, 59.04, 49.06, 42.68, 29.76. HRMS (ESI-MS): m/z [M+H^+^] calculated for C_13_ H_18_ N_3_ O^+^: 232.1444, found 232.1449; C_13_H_17_N_3_O (231.30).

##### 8-(4-(4-Fluorophenyl)-4-oxobutyl)-1-phenyl-1,3,8-triazaspiro[4.5]decan-4-one (6)^[26]^

**6** (1.40 g, 6.0 mmol, 1 eq), 4-chloro-1-(4-fluorophenyl)butan-1-one (1.79 g, 9.0 mmol, 1.5 eq), NaI (1.35 g, 9.0 mmol, 1.5 eq) and triethylamine (0.91 g, 9.0 mmol, 1.5 eq) were suspended in MeCN (35 mL) and refluxed under argon atmosphere overnight. After cooling to room temperature the solvent was evaporated. The resulting crude residue was dissolved in DCM (35 mL) and washed with aqueous KOH (20%, 20 mL). The organic layer was dried over Na_2_SO_4_ and concentrated under reduced pressure. The residue was dissolved in DCM (20 mL) and the solution was added dropwise to cold hexane (120 mL). The gray precipitate was filtered and dried in vacuo to give **7** (1.31 g, 55%). ^1^H NMR (400 MHz, DMSO-d_6_) δ 8.58 (s, 1H), 8.10 – 8.01 (m, 2H), 7.39 – 7.29 (m, 2H), 7.24 – 7.10 (m, 3H), 6.79 – 6.68 (m, 3H), 4.54 (s, 2H), 3.02 (t, *J* = 6.9 Hz, 2H), 2.75 – 2.59 (m, 4H), 2.48 – 2.33 (m, 5H), 1.88 – 1.74 (m, 2H), 1.56 – 1.46 (m, 2H). ^13^C NMR (101 MHz, DMSO-*d*_6_) δ 199.10, 176.67, 164.8 (d, *J* = 251.0 Hz), 143.82, 131.29 (d, *J* = 9.4 Hz), 129.42, 118.05, 116.10 (d, J = 21.7 Hz), 114.71, 114.39, 59.09, 58.74, 57.63, 49.68, 36.39, 28.86, 22.05. HRMS (ESI-MS): m/z [M+H^+^] calculated for C_23_ H_27_ FN_3_ O_2_ ^+^: 396.2082, found 396.2087; C_23_H_26_FN_3_O_2_ (395.48).

##### tert-Butyl (4-(2-hydroxyethyl)phenyl)carbamate (8)^[30]^

Di-tert-butyl dicarbonate (1.75 g, 8.0 mmol, 1.1 eq) was dissolved in EtOAc (20 mL) and added dropwise to a suspension of 2-(4-aminophenyl)ethan-1-ol (**7**, 1.00 g, 7.3 mmol, 1 eq) in EtOAc (10 mL). After the reaction was stirred at room temperature overnight the solvent was evaporated. The resulting crude was purified by column chromatography (DCM/MeOH 95/5) to afford **8** (1.48 g, 86%) as a white foam. ^1^H NMR (300 MHz, CDCl_3_) δ 7.33 – 7.25 (m, 2H), 7.17 – 7.07 (m, 2H), 6.59 (bs, 1H), 3.79 (t, *J* = 6.6 Hz, 2H), 2.80 (t, *J* = 6.6 Hz, 2H), 1.51 (s, 9H). ^13^C NMR (75 MHz, CDCl_3_) δ 152.97, 136.80, 133.13, 129.54, 118.99, 80.50, 63.71, 38.49, 28.37. HRMS (ESI-MS): m/z [M+H^+^] calculated for C_13_ H_20_ NO_3_ ^+^: 238.1438, found 238.1437; C_13_ H_19_ NO_3_ (237.30).

##### tert-Butyl (4-(2-bromoethyl)phenyl)carbamate (9)^[30]^

To a cooled solution of **8** (1.48 g, 6.3 mmol, 1 eq) in DCM (30 mL) triphenyl phosphine (2.50 g, 9.5 mmol, 1.5 eq) and N-bromosuccinimide (1.71 g, 9.5 mmol, 1.5 eq) were added. The mixture was stirred for 2 h while maintaining the temperature at 0°C. Then, the solvent was evaporated and the resulting residue was purified by column chromatography (DCM/MeOH 99/1) to afford **9** (1.68 g, 88%) as a white semi solid. ^1^H NMR (300 MHz, CDCl_3_) δ 7.36 – 7.25 (m, 2H), 7.17 – 7.08 (m, 2H), 6.47 (bs, 1H), 3.52 (t, *J* = 7.6 Hz, 2H), 3.10 (t, J = 7.6 Hz, 2H), 1.51 (s, 9H). ^13^C NMR (101 MHz, CDCl _3_) δ 152.76, 137.17, 133.55, 129.84, 129.23, 124.88, 118.78, 38.77, 33.12, 28.35. HRMS (ESI-MS): m/z [M+H^+^] calculated for C_13_ H_19_ BrNO_2_ ^+^: 300.0594, found 300.0593; C_13_ H_18_ BrNO_2_ (300.20).

##### tert-Butyl (4-(2-(8-(4-(4-fluorophenyl)-4-oxobutyl)-4-oxo-1-phenyl-1,3,8-triazaspiro[4.5]decan-3-yl)ethyl)phenyl)carbamate (10)^[26]^

A mixture of **7** (0.79 g, 2.0 mmol, 1 eq), potassium hydroxide (0.056 g, 1.0 mmol, 0.5 eq), potassium carbonate (1.09 g, 8.0 mmol, 4 eq) and tetrabutylammonium bisulfate (0.20 g, 0.6 mmol, 0.3 eq) was suspended in anhydrous toluene (40 mL) and stirred for 30 min under argon atmosphere. Then, a solution of **9** (1.21 g, 4.0 mmol, 2 eq) in anhydrous toluene (25 mL) was added dropwise over 30 min. The reaction was stirred at 90°C for 2 days. After that, the reaction was allowed to cool to room temperature and the solvent was evaporated. The resulting crude residue was dissolved in DCM (40 mL) and washed with aqueous KOH (20%, 20 mL). The organic layer was dried over Na_2_SO_4_ and concentrated under reduced pressure. The residue was purified by column chromatography (DCM/MeOH 95/5) to afford **10** (0.41 g, 35%) as a white foam. ^1^H NMR (400 MHz, CDCl_3_) δ 8.03 – 7.96 (m, 2H), 7.30 – 7.21 (m, 4H), 7.16 – 7.08 (m, 4H), 6.90 – 6.80 (m, 3H), 6.48 (s, 1H), 4.52 (s, 2H), 3.63 (t, J = 7.1 Hz, 2H), 3.15 – 2.49 (m, 12H), 2.09 – 1.96 (m, 2H), 1.54 – 1.49 (m, 2H), 1.48 (s, 9H). ^13^C NMR (75 MHz, CDCl_3_) δ 198.38, 174.13, 167.37, 163.99, 152.84, 142.80, 137.16, 133.51, 133.47, 132.49, 130.79, 130.67, 129.30, 129.21, 119.00, 118.86, 115.82, 115.54, 115.37, 80.48, 63.80, 60.28, 57.47, 49.43, 42.21, 36.33, 33.02, 28.92, 28.36, 21.32. HRMS (ESI-MS): m/z [M+H^+^] calculated for C_36_ H_44_ FN_4_ O_4_^+^: 615.3341, found 615.3347; C_36_ H_43_ FN_4_ O_4_ (614.76).

##### 3-(4-Aminophenethyl)-8-(4-(4-fluorophenyl)-4-oxobutyl)-1-phenyl-1,3,8-triazaspiro[4.5]decan-4-one (11)^[26]^

**10** (0.41 g, 0.7 mmol) was dissolved in DCM (30 mL) and TFA (5 mL) was added. The mixture was stirred at room temperature overnight. After the reaction was complete, as indicated by TLC, the solution was basified with aqueous KOH (20%). The organic phase was separated, dried over Na_2_SO_4_ and concentrated under reduced pressure. The residue was purified by column chromatography to yield **11** (0.28 g, 83%). as a yellow oil. ^1^H NMR (400 MHz, CDCl_3_) δ 8.06 – 7.98 (m, 2H), 7.28 – 7.19 (m, 2H), 7.16 – 7.10 (m, 2H), 7.03 – 6.96 (m, 2H), 6.87 – 6.79 (m, 3H), 6.67 – 6.57 (m, 2H), 4.52 (s, 2H), 3.66 – 3.54 (m, 2H), 3.04 (t, J = 7.1 Hz, 2H), 2.97 – 2.53 (m, 10H), 2.08 – 1.97 (m, 2H), 1.55 (d, J = 14.1 Hz, 2H).^13^C NMR (101 MHz, CDCl_3_) δ 198.01, 173.84, 165.72 (d, J = 254.5 Hz), 145.17, 142.68, 133.39, 130.71 (d, J = 9.2 Hz), 129.53, 129.35, 127.62, 119.02, 115.7 (d, J = 21.9 Hz), 115.31, 115.14, 63.75, 59.96, 57.22, 49.26, 42.31, 36.10, 32.82, 28.56, 20.89. HRMS (ESI-MS): m/z [M+H^+^] calculated for C_31_ H_36_ FN_4_ O_2_ ^+^: 515.2817, found 515.2820; C_31_ H_35_ FN_4_ O_2_ (514.65).

##### tert-Butyl (3-(2-(2-(3-aminopropoxy)ethoxy)ethoxy)propyl)carbamate (12)^[34]^

To a solution of 3,3’-((oxybis(ethane-2,1-diyl))bis(oxy))bis(propan-1-amine) (1.76 g, 8.0 mmol, 4 eq) in DCM (35 mL) a solution of di-tert-butyl dicarbonate (0.44 g, 2.0 mmol, 1 eq) in DCM (20 mL) was added dropwise over 30 min. The mixture was stirred at room temperature for 16 h. The solvent was evaporated and the residue was purified by column chromatography (DCM/MeOH 9/1 + 0.1% NH_3_) to afford **12** (0.61 g, 95%) as a yellow oil. ^1^H NMR (300 MHz, CDCl_3_) δ 3.89 (s, 2H), 3.67 – 3.56 (m, 10H), 3.56 – 3.49 (m, 2H), 3.30 – 3.16 (m, 2H), 3.03 (t, J = 6.0 Hz, 2H), 1.97 – 1.84 (m, 2H), 1.82 – 1.70 (m, 2H), 1.42 (s, 9H). ^13^C NMR (75 MHz, CDCl_3_) δ 160.28, 81.30, 70.56, 70.38, 70.13, 70.08, 69.76, 69.40, 39.74, 38.41, 29.70, 28.48. HRMS (ESI-MS): m/z [H+M^+^] calculated for C_15_ H_33_ N_2_ O_5_ ^+^: 321.2384, found 321.2390; C_15_H_32_N_2_O_5_ (320.43).

##### 4-((tert-Butoxycarbonyl)amino)butanoic acid (13)^[35]^

To a mixture of 4-aminobutryric acid (0.30 g, 2.9 mmol, 1 eq) and NaOH (0.12 g, 2.9 mmol, 1 eq) in dioxane/water (1/1, 30 mL) a solution of Boc_2_O (0.70 g, 3.2 mmol, 1.1 eq) was added in dioxane (30 mL). After the reaction was stirred at room temperature overnight aqueous HCl (1 N) was added to set pH < 1. The mixture was extracted three times with EtOAc (3 x 30 mL) and the organic layer was dried over Na_2_SO_4_. The solvent was removed under reduced pressure and the crude product was purified by column chromatography (DCM/MeOH 95/5 to 9/1) to afford **13** as a colorless oil (0.38 g, 64%). ^1^H NMR (400 MHz, CDCl_3_) δ 3.17 (t, J = 6.7 Hz, 2H), 2.39 (t, J = 7.2 Hz, 2H), 1.81 (p, J = 7.0 Hz, 2H), 1.44 (s, 9H). ^13^C NMR (101 MHz, CDCl_3_) δ 178.05, 156.20, 79.53, 39.93, 31.27, 28.37, 25.17. HRMS (ESI-MS): m/z [M+H^+^] calculated for C_9_ H_18_ NO_4_ ^+^: 204.1230, found 204.1231; C_9_ H_17_ NO_4_ (203.24).

##### tert-Butyl (18-((4-(2-(8-(4-(4-fluorophenyl)-4-oxobutyl)-4-oxo-1-phenyl-1,3,8-triazaspiro[4.5]decan-3-yl)ethyl)phenyl)amino)-15,18-dioxo-4,7,10-trioxa-14-azaoctadecyl)carbamate (14)^[26]^

To a solution of **11** (0.28 g, 0.54 mmol, 1 eq) in DMF (30 mL) succinic anhydride (0.054 g, 0.54 mmol, 1 eq) was added and the reaction was stirred at room temperature overnight. After the reaction was complete, as indicated by TLC, HATU (0.31 g, 0.82 mmol, 1.5 eq), DIPEA (0.21 g, 1.62 mmol, 3 eq) and **12** (0.17 g, 0.54 mmol, 1 eq) were added. Then, the mixture was stirred for 10 h at room temperature. The solvent was evaporated and the residue was purified by column chromatography (DCM/MeOH 95/5). **14** (0.25 g, 47%) was obtained as a brown oil. ^1^H NMR (400 MHz, CDCl_3_) δ 8.02 – 7.96 (m, 2H), 7.45 (d, *J* = 8.3 Hz, 2H), 7.30 (t, *J* = 8.0 Hz, 2H), 7.18 – 7.10 (m, 4H), 6.97 (d, *J* = 8.3 Hz, 2H), 6.93 – 6.83 (m, 2H), 5.00 (s, 1H), 4.59 (s, *J* = 22.1 Hz, 2H), 4.40 (s, 1H), 3.76 – 3.64 (m, 4H), 3.64 – 3.59 (m, 4H), 3.59 – 3.42 (m, 10H), 3.35 (dd, *J* = 11.8, 5.8 Hz, 2H), 3.28 – 3.08 (m, 8H), 2.97 – 2.88 (m, 2H), 2.70 – 2.63 (m, 2H), 2.62 – 2.54 (m, 2H), 2.35 – 2.23 (m, 2H), 1.80 – 1.67 (m, 4H), 1.52 (d, *J* = 14.1 Hz, 2H), 1.42 (s, 9H). ^13^C NMR (101 MHz, CDCl_3_) δ 196.67, 172.80, 171.00, 165.98 (d, J = 255.5 Hz), 156.19, 141.80, 137.20, 132.79, 132.75, 130.78 (d, *J* = 9.4 Hz)129.75, 129.18, 120.24, 119.62, 115.89 (d, *J* = 21.9 Hz), 114.80, 79.10, 70.49, 70.17, 70.04, 69.42, 63.47, 58.48, 56.57, 48.67, 41.58, 38.42, 35.51, 33.02, 32.92, 31.60, 29.70, 28.64, 28.45, 26.81, 18.21. HRMS (ESI-MS): m/z [H+M^+^] calculated for C_50_ H_70_ FN_6_ O_6_ ^+^: 917.5183, found 917.5194; C_50_ H_69_ FN_6_ O_6_ (917.13)

##### N^1^-(3-(2-(2-(3-Aminopropoxy)ethoxy)ethoxy)propyl)-N^4^-(4-(2-(8-(4-(4-fluorophenyl)-4-oxobutyl)-4-oxo-1-phenyl-1,3,8-triazaspiro[4.5]decan-3-yl)ethyl)phenyl)succinamide trihydrotrifluoracetate (15)^[26]^

**14** (0.05 g, 0.07 mmol) was dissolved in DCM (30 mL) and TFA (5 mL) was added. The mixture was stirred at room temperature overnight. After the reaction was complete, as indicated by TLC, the solvent was evaporated. The resulting crude product was purified by preparative HPLC. **15** (18.1 mg, 22%) was obtained as a yellow oil. ^1^H NMR (400 MHz, D_2_O) δ 8.04 – 7.95 (m, 2H), 7.43 – 7.33 (m, 4H), 7.32 – 7.19 (m, 4H), 7.13 – 6.92 (m, 3H), 4.63 (s, 2H), 3.77 – 3.68 (m, 2H), 3.66 – 3.55 (m, 8H), 3.54 – 3.50 (m, 2H), 3.50 – 3.03 (m, 14H), 3.00 – 2.88 (m, 2H), 2.75 – 2.58 (m, 2H), 2.58 – 2.48 (m, 2H), 2.48 – 2.33 (m, 2H), 2.12 – 1.99 (m, 2H), 1.99 – 1.88 (m, 2H), 1.85 – 1.62 (m, 4H). ^13^C NMR (101 MHz, D_2_O) δ 200.77, 174.41, 173.34, 172.89, 165.98 (d, *J* = 255.2 Hz), 141.75, 135.86, 135.02, 132.50, 131.05 (d, *J* = 9.8 Hz), 129.75, 129.66, 122.13, 121.36, 118.72, 117.86, 115.86 (d, *J* = 22.2 Hz), 69.52, 69.43, 69.28, 68.31, 63.55, 59.29, 56.08, 48.88, 41.60, 37.69, 36.29, 34.90, 32.08, 31.91, 31.07, 28.31, 27.17, 26.52, 18.14. HRMS (ESI-MS): m/z [M+H^+^] calculated for C_45_ H_62_ FN_6_ O_7_ ^+^: 817.4659, found 817.4654; C_45_H_61_FN_6_O_7_ x C_6_H_3_F_9_O_6_ (817.02 + 342.07)

##### Diazapentacosan-25-yl)-3-methyl-5-sulfonato-3-(3-sulfonatopropyl)indolin-2-ylidene)prop-1-en-1-yl)-1-(2-methoxyethyl)-3-methyl-3-(3-sulfonatopropyl)-3H-indol-1-ium-5-sulfonate (16)

**15** (0.333 mg, 0.288 μmol, 1.5 eq) was dissolved in DMF (30 μL). NEt_3_ (0.20 mg, 1.92 μmol, 10 eq) and Dyomics Dye 549-P1 NHS ester (0.2 mg, 0.192 μmol, 1 eq) in DMF (60 μL) was added and the reaction was shaken for 2.5 h in the dark at room temperature. The reaction was quenched with 10% aqueous TFA (20 μL) and the crude product was purified by preparative HPLC (method B). **15** (0.3 mg, 89%) was obtained as a pink solid. Anal. RP-HPLC (254 nm) (method B): 99% (t_R_ = 7.22 min, k = 1.25). HRMS (ESI-MS): m/z [M+3H^3+^] calculated for C_81_ H_110_ FN_8_ O_21_ S4^3+^: 559.2212, found 559.2226; C_81_ H_107_ FN_8_ O_21_ S_4_ x N_3_H_9_ (1673.00 + 54.12).

##### 2-(6-(Dimethylamino)-3-(dimethyliminio)-3H-xanthen-9-yl)-5-((18-((4-(2-(8-(4-(4-fluorophenyl)-4-oxobutyl)-4-oxo-1-phenyl-1,3,8-triazaspiro[4.5]decan-3-yl)ethyl)phenyl)amino)-15,18-dioxo-4,7,10-trioxa-14-azaoctadecyl)carbamoyl)benzoate dihydrotrifluoroacetate (17)

**15** (1.8 mg, 1.56 μmol, 1.2 eq) was dissolved in DMF (30 μL). NEt_3_ (1.1 mg, 10.4 μmol, 10 eq) and 5-carboxytetramethylrhodamine succinimidyl ester (5-TAMRA NHS ester) (0.65 mg, 1.28 μmol, 1 eq) in DMF (60 μL) were added and the reaction was shaken for 2.5 h in the dark at room temperature. The reaction was quenched with 10% aqueous TFA (20 μL) and the crude product was purified by preparative HPLC (method A). A pink solid was obtained for **17** (1.58 mg, 1.08 μmol, 84%). Anal. RP-HPLC (254 nm) (method A): 99% (t_R_ = 13.38 min, k = 3.17) HRMS (ESI-MS): m/z [M+H^+^] calculated for C_70_ H_82_ FN_8_ O_11_ ^+^: 1229.6082, found 1229.6092; C_70_ H_81_ FN_8_ O_4_ x C_4_ H_2_ F_6_ O_4_ (1229.46 + 228.05).

##### tert-Butyl (4-((4-(2-(8-(4-(4-fluorophenyl)-4-oxobutyl)-4-oxo-1-phenyl-1,3,8-triazaspiro[4.5]decan-3-yl)ethyl)phenyl)amino)-4-oxobutyl)carbamate (18)

A mixture of **13** (23 mg, 0.11 mmol, 1.1 eq) and HATU (57 mg, 0.15 mmol, 1.5 eq) in DMF (15 mL) was stirred at 0°C for 10 min. Then, DIPEA (40 mg, 0.3 mmol, 3 eq) and **11** (50 mg, 0.1 mmol, 1 eq) were added slowly and the reaction was stirred at room temperature overnight. After the solvent was removed under reduced pressure the residue was dissolved in DCM (10 mL) and washed three times with aqueous KOH (20%, 3 x 10 mL). The organic layer was dried over Na_2_SO_4_ and the solution was concentrated in vacuo. The crude product was purified by column chromatography (DCM/MeOH 99/1 to 95/5) to give **18** as a yellow oil (63 mg, 90%). ^1^H NMR (400 MHz, CDCl_3_) δ 9.01 (d, *J* = 17.7 Hz, 1H), 8.17 – 8.08 (m, 2H), 7.67 – 7.62 (m, 2H), 7.35 (dd, J = 8.4, 7.6 Hz, 2H), 7.28 – 7.20 (m, 4H), 7.04 – 6.90 (m, 3H), 5.11 (t, *J* = 5.7 Hz, 1H), 4.66 (s, *J* = 7.6 Hz, 2H), 3.77 (t, *J* = 7.1 Hz, 2H), 3.32 (d, *J* = 5.3 Hz, 2H), 3.20 – 2.63 (m, 12H), 2.50 – 2.46 (m, 2H), 2.10 (d, *J* = 26.2 Hz, 2H), 2.01 – 1.93 (m, 3H), 1.66 (d, *J* = 14.0 Hz, 2H), 1.55 (s, *J* = 1.7 Hz, 9H). ^13^C NMR (101 MHz, CDCl_3_) δ 198.23, 173.98, 171.32, 165.7 (d, *J* = 254.5 Hz), 157.15, 142.73, 137.28, 136.83, 134.28, 133.41, 133.31, 130.72 (d, *J* = 9.2 Hz), 129.40, 129.32, 129.09, 120.19, 120.06, 119.16, 115.69 (d, *J* = 21.8 Hz), 115.49, 79.68, 63.77, 63.58, 60.10, 57.35, 53.47, 49.37, 42.09, 39.39, 38.66, 36.20, 34.56, 33.10, 28.41, 27.09. HRMS (ESI-MS): m/z [M+H^+^] calculated for C_40_ H_51_ FN_5_ O_5_ ^+^: 700.3869, found 700.3875; C_40_ H_50_ FN_5_ O_5_ (699.87).

##### 4-Amino-N-(4-(2-(8-(4-(4-fluorophenyl)-4-oxobutyl)-4-oxo-1-phenyl-1,3,8-triazaspiro[4.5]decan-3-yl)ethyl)phenyl)butanamide trihydrotrifluoracetate (19)

**15** (63 mg, 0.09 mmol) was dissolved in DCM (30 mL) and TFA (5 mL) was added. The mixture was stirred at room temperature overnight. After the reaction was complete, as indicated by TLC, the solvent was evaporated. The resulting crude product was purified by preparative HPLC. **19** (32.1 mg, 60%) was obtained as a white foam-like solid. ^1^H NMR (400 MHz, CD_3_ OD) δ 8.10 – 8.04 (m, 2H), 7.47 (d, *J* = 8.4 Hz, 2H), 7.28 – 7.18 (m, 6H), 6.97 – 6.86 (m, 3H), 4.65 (s, 2H), 3.79 – 3.64 (m, 4H), 3.54 (d, *J* = 8.8 Hz, 2H), 3.25 – 3.14 (m, 4H), 3.06 – 2.92 (m, 4H), 2.76 – 2.63 (m, 2H), 2.51 (t, *J* = 7.0 Hz, 2H), 2.18 – 2.07 (m, 2H), 2.05 – 1.94 (m, 2H), 1.74 (d, *J* = 14.7 Hz, 2H). ^13^C NMR (101 MHz, CD_3_OD) δ 197.32, 172.82, 165.95 (d, *J* = 253.4 Hz)171.37, 142.18, 136.94, 134.07, 133.07, 130.64 (d, *J* = 9.5 Hz) 129.10, 129.06, 120.34, 120.18, 116.90, 115.29 (d, J = 22.2 Hz) 63.45, 58.44, 56.25, 49.04, 41.40, 39.00, 34.53, 32.96, 32.27, 27.21, 22.86, 18.17. HRMS (ESI-MS): m/z [M+H^+^] calculated for C H FN O ^+^: 600.3344, found 600.3345; C_35_ H_42_ FN_5_ O_3_ x C_6_ H_3_ F_9_ O_6_ (599.75 + 342.07)

##### 2-(6-(Dimethylamino)-3-(dimethyliminio)-3H-xanthen-9-yl)-5-((4-((4-(2-(8-(4-(4-fluorophenyl)-4-oxobutyl)-4-oxo-1-phenyl-1,3,8-triazaspiro[4.5]decan-3-yl)ethyl)phenyl)amino)-4-oxobutyl)carbamoyl)benzoate dihydrotrifluoracetate (20)

**19** (1.8 mg, 1.92 μmol, 1.5 eq) was dissolved in DMF (30 μL). NEt_3_ (1.1 mg, 10.4 μmol, 10 eq) and 5-carboxytetramethylrhodamine succinimidyl ester (5-TAMRA NHS ester) (0.65 mg, 1.28 μmol, 1 eq) in DMF (60 μL) were added and the reaction was shaken for 2.5 h in the dark at room temperature. The reaction was quenched with 10% aqueous TFA (20 μL) and the crude product was purified by preparative HPLC (method A). A pink solid was obtained for **20** (0.84 mg, 0.75 μmol, 59%). Anal. RP-HPLC (254 nm) (method A): 99% (*t*_R_ = 13.20 min, k = 3.15). HRMS (ESI-MS): m/z [M+H^+^] calculated for C_60_H_63_FN_7_O_7_ ^+^: 1012.4768, found 1012.4764; C_60_H_62_FN_7_O_7_ x C_4_H_2_F_6_O_4_ (1012.19 + 228.05).

### Radioligand competition binding experiments at the dopamine receptors

Cell homogenates containing the D_2long_R, D_3_R, and D_4.4_R were kindly provided by Dr. Lisa Forster, University of Regensburg. Homogenates containing the D_1_R and D_5_R were prepared and radioligand binding experiments with cell homogenates were performed as previously described with minor modifications.^[36,37]^ For radioligand competition binding assays homogenates were incubated in BB at a final concentration of 0.3 μg (D_1_R), 0.3 μg (D_2long_R), 0.7 μg (D_3_R), 0.5-1.0 μg (D_4.4_R), or 0.4 μg (D_5_R) protein/well. [^3^H]SCH-23390 (D_1_R (*K*_d_ = 0.23 nM) and D_5_R (*K*_d_ = 0.20 nM)) was added in final concentrations of 1.0 nM (D_1_R) and 1.0 nM (D_5_R). [^3^H]N-methylspiperone (D_2long_R (*K*_d_ = 0.0149 nM), D_3_R (*K*_d_ = 0.0258 nM), D_4.4_R (*K*_d_ = 0.078 nM)) was added in final concentrations of 0.05 nM (D_2long_R, D_3_R) or 0.1 nM (D_4.4_R). Non labelled compounds were added in increasing concentrations for the displacement of the radioligands. After incubation of 60 min (D_2long_R, D_3_R, and D_4.4_R) or 120 min (D_1_R and D_5_R) at room temperature, bound radioligand was separated from free radioligand through PEI-coated GF/C filters using a Brandel harvester (Brandel Inc., Unterföhring, Germany), filters were transferred to (flexible) 1450-401 96-well sample plates (PerkinElmer, Rodgau, Germany) and after incubation with scintillation cocktail (Rotiszint eco plus, Carl Roth, Karlsruhe, Germany) for at least 3 h, radioactivity was measured using a MicroBeta2 plate counter (PerkinElmer, Waltham, MA, USA). Competition binding curves were fitted using a four-parameter fit (“log(agonist) vs. response-variable slope”). Calculations of pK_i_ values with SEM and graphical presentations were conducted with GraphPad Prism 9 software (San Diego, CA, USA).

### G_o1_ heterotrimer dissociation assay

HEK293A cells (Thermo Fisher) were transiently transfected with the G_o1_ BRET sensor, G_o1_-CASE (https://www.addgene.org/168123/),^[32]^ along with either D_2long_ R, D_3_ R or D_4_ R using polyethyleneimine (PEI). Per well of a 96-well plate, 100 μl of freshly resuspended cells were incubated with 100 ng total DNA (50 ng receptor and 50 ng G_o1_-CASE) mixed with 0.3 μl PEI solution (1 mg/ml) in 10 μl Opti-MEM (Thermo Fisher), seeded onto poly-D-lysine (PDL)-pre-coated white, F-bottom 96-well plates (Brand GmbH) and cultivated at 37°C, 5% CO_2_ in penicillin (100 U/ml)/streptomycin (0.1 mg/ml)-, 2 mM L-glutamin- and 10% fetal calf serum (FCS)-supplemented Dulbecco’s modified Eagle’s medium (DMEM; Thermo Fisher). 48 hours after transfection, cells were washed with Hank’s Buffered Salt Solution (HBSS) and incubated with a 1/1000 furimazine (Promega; cat no. N1663) dilution in HBSS for 2 minutes. Next, the baseline BRET ratio was recorded in three consecutive reads, cells were stimulated with serial dilutions of **20** or vehicle control, and BRET was recorded for another 15 reads. For experiments in antagonist mode, serial dilutions of **20** were added together with furimazine before the experiment and 1 μM dopamine or vehicle control was added after the first three baseline recordings. All experiments were conducted using a ClarioStar Plus Plate reader (BMG Labtech) with a cycle time of 120 seconds, 0.3 seconds integration time and a focal height of 10 mm. Monochromators were used to collect the NanoLuc emission intensity between 430 and 510 nm and cpVenus emission between 500 and 560 nm. BRET ratios were defined as acceptor emission/donor emission. The basal BRET ratio before ligand stimulation (Ratio_basal_) was defined as the average of all three baseline BRET values. Ligand-induced ΔBRET was calculated for each well as a percent over basal ([(Ratio_stim_ − Ratio_basal_)/Ratio_basal_] × 100). To correct for non-pharmacological effects, the average ΔBRET of vehicle control was subtracted.

### Fluorescence Properties

Excitation and emission spectra of **16, 17**, and **20** were recorded in PBS (137 mM NaCl, 2.7 mM KCl, 10 mM Na_2_HPO_4_, 1.8 mM KH_2_PO_4_, pH 7.4) containing 1% BSA (Sigma-Aldrich, Munich, Germany) using a Cary Eclipse spectrofluorometer (Varian Inc., Mulgrave, Victoria, Australia) at 22 °C, in acryl cuvettes (10 ×10 mm, Sarstedt, Nümbrecht, Germany). The slit adjustments (excitation/emission) were 5/10 nm for excitation spectra and 10/5 nm for emission spectra. Net spectra were calculated by subtracting the respective vehicle reference spectrum, and corrected emission spectra were calculated by multiplying the net emission spectra with the respective lamp correction spectrum. The quantum yields of **16, 17**, and **20** were determined according to previously described procedures^[33,38]^ with minor modifications using a Cary Eclipse spectrofluorometer (Varian Inc., Mulgrave, Victoria, Australia) at 22 °C, using acryl cuvettes (10 ×10 mm, Sarstedt, Nümbrecht, Germany) and cresyl violet perchlorate (Biomol GmbH −Life Science Shop, Hamburg, Germany) as a red fluorescent standard. Absorption spectra were recorded by UV/Vis spectroscopy (350−850 nm, scan rate: 300 nm/min, slits: fixed 2 nm) at a concentration of 2 μM for cresyl violet (in EtOH, λ_abs_,_max_ = 575 nm) and **16, 17**, and **20** (in PBS + 1% BSA, λ_abs_,_max_ = 559-562 nm). The excitation wavelength for the emission spectra was 550 nm (**16**) or 545 nm (**17, 20**). The emission wavelength collected during the excitation scan was 590 nm (**16**) or 610 nm (**17, 20**). The quantum yields were calculated for three different slit adjustments (exc./em.): 5/5, 10/5, and 10/10 nm. The means of the quantum yields, absorption and emission maxima are presented in Table 3.

### Live cell confocal microscopy at the D_2long_R

Confocal images were recorded with kind assistance from Manel Bosch (Universitat de Barcelona). HEK293T cells were transfected based on previously described procedures with the D_2long_R-GFP_2_ (C-terminally tagged) with minor modifications.^[39–41]^ Cells were seeded in 35 mm wells containing 1.5 cover slips. 48 h after transfection medium was changed to OptiMem media (Gibco) supplemented with 10 mM HEPES. Imaging was performed using a Zeiss LSM880 Laser Scanning Confocal Microscope equipped with a “Plan-Apochromat” 40x/1,3 Oil DIC M27 objective and a photomultiplier tube (PTM) detector. For excitation of GFP_2_ an argon Laser with a wavelength of 488 nm was used. Fluorescence was detected within an emission window of 505-605 nm. For excitation of **20** an DPSS laser with a wavelength of 561 nm was used. Fluorescence was detected within an emission window of 569-669 nm. Image size was set to 512 x 512 pixels. After adjusting the focus, time-lapse images were recorded. **20** was added in a final concentration of 50 nM.

### Molecular Brightness

HEK-293AD cells were seeded in 8-well Ibidi μ-slides with a density of 25,000 cells per well and transfected with 2 μg hD_2_R-mNeonGreen after 24 h using JetPrime transfection reagent according to manufacturer’s protocol. After further 24 h cells were washed and imaged in FRET-buffer (144 mM NaCl, 5.4 mM KCl, 1 mM MgCl_2_, 2 mM CaCl_2_, 10 mM HEPES) on a confocal laser scanning microscope, Leica SP8, with a white-light laser at wavelengths of 488 and 552 nm, and laser power of 5%. All measurements were conducted with an HC PLAP CS2 40 × 1.3 numerical aperture (NA) oil immersion objective (Leica). Movies were acquired at 1.3 seconds per frame for 100 frames with two hybrid detectors in the range of 498 to 547 nm and 562 to 612 nm respectively, in a line sequential, counting mode. Molecular brightness ε and number of molecules N are calculated from the average (k) of the photon counts collected in a pixel and its variance (σ) according to the formulas ε = σ^2^/k-1 and N= k^2^/ σ^2^. ImageJ was used to extract molecular brightness and fluorescence intensity values, number of emitters was calculated with Word Excel and obtained values were plotted and fitted with Prism v. 9.5.1.

## Notes

The authors declare no competing financial interest.

## Author contribution

The authors contributed as follows: M.N.: synthesis and analytics of the compounds, radioligand binding studies, fluorescence properties and manuscript writing. D.M.: radioligand binding studies, preparation of mNeonGreen constructs. N.R.: synthesis and radioligand binding studies. H.S.: G_o1_ heterotrimer dissociation assay, manuscript writing. A.S.: Molecular Brightness studies, manuscript writing. N.K.: Molecular Brightness studies. I.R.-R.: Cell culture for confocal microscopy. G.N.: Cell culture for confocal microscopy, conception. R.F.: conception, providing infrastructure. P.K.: conception, providing infrastructure. P.A.: conception, manuscript writing, providing infrastructure. S.P.: conception, project administration, manuscript writing, confocal microscopy, providing infrastructure. The data were analyzed and discussed by all authors. All authors have given approval to the final version of the manuscript.

## Supporting Information

The Supporting Information is available free of charge at link DOI: XXX/number

Chemical purity and stability (Figure S1-S4), Dopamine-induced G_o1_ activation (Figure S5), NMR spectra (Figure S6-S37), Structures of the fluorescent ligands **16, 17** and **20** (Figure S38-S40) (PDF).

## Acknowledgements

We thank Manel Bosch (Universitat de Barcelona) for expert technical assistance in confocal microscopy. We thank Dr. Lisa Forster for providing cell homogenates for the D_2long_R, D_3_R and D_4_R. We thank Prof. Dr. Sigurd Elz for providing infrastructure. S.P. was supported by the Fonds der Chemischen Industrie (No. 661688) and University of Regensburg (Academic Research Sabbatical Program). Financial support by the graduate school “Receptor Dynamics: Emerging Paradigms for Novel Drugs (K-BM-2013-247)” of the Elite Network of Bavaria (ENB) for S.P., N.R., M.N. and H.S. is gratefully acknowledged. We would like to thank the Deutsche Forschungsgemeinschaft (DFG) for support through project 421152132 SFB1423 subproject C03 (P.A.) and SFB 1470 subproject A01 (P.A.). R.F., as PI, was funded by Spanish MCIN/AEI/10.13039/501100011033 (grant PID2021-126600OB-I00) and by the European Union Next Generation EU/PRTR (ERDF A way of making Europe).

