## Supporting Information for "Fluorescent Tools for Imaging and Ligand Screening of Dopamine D_2_-Like Receptors"

### Contents


1. Chemical purity and stability ........ S3
2. Dopamine-induced $G_{o1}$ activation ........ S5
3. NMR spectra ........ S6
4. Structures of the fluorescent ligands **16**, **17** and **20** ........ S22

### 1. Chemical purity and stability

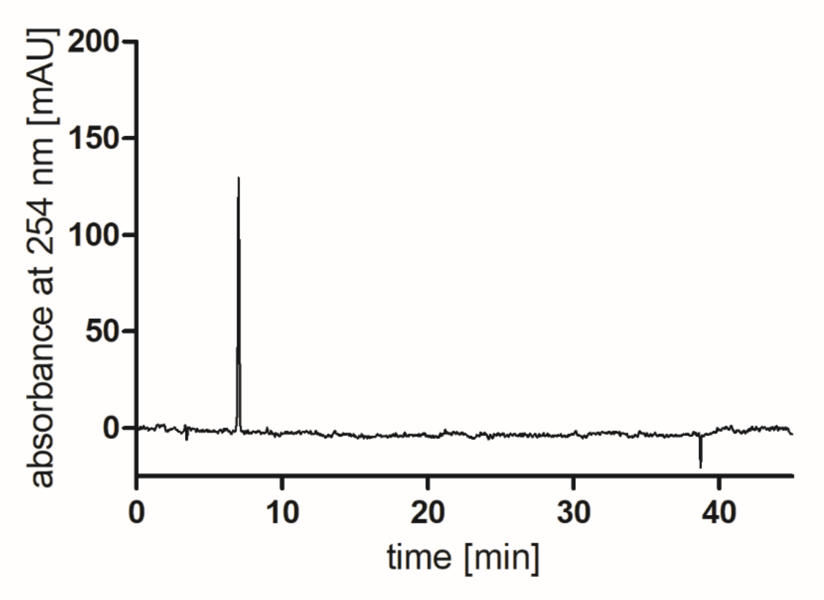


**Figure S1.** RP-HPLC analysis (purity control) of compound **16** (>99%, 254nm).

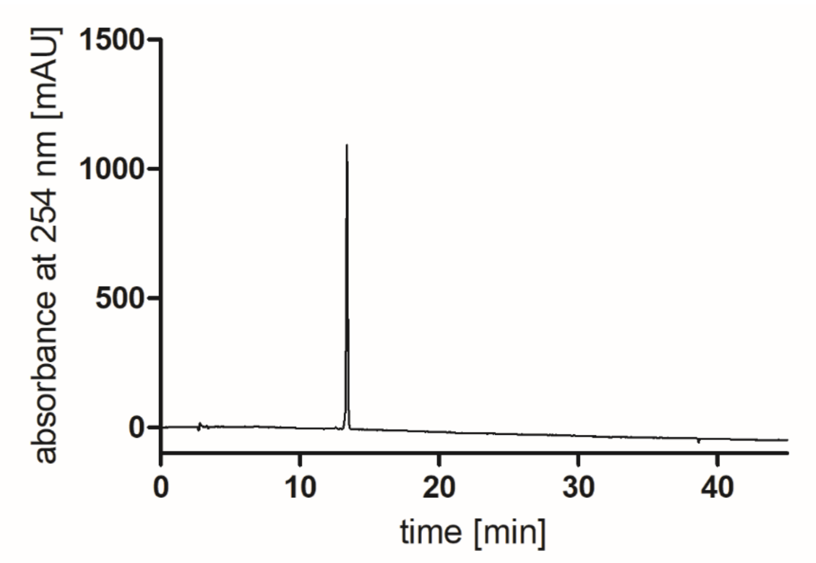


**Figure S2.** RP-HPLC analysis (purity control) of compound **17** (>99%, 254nm).

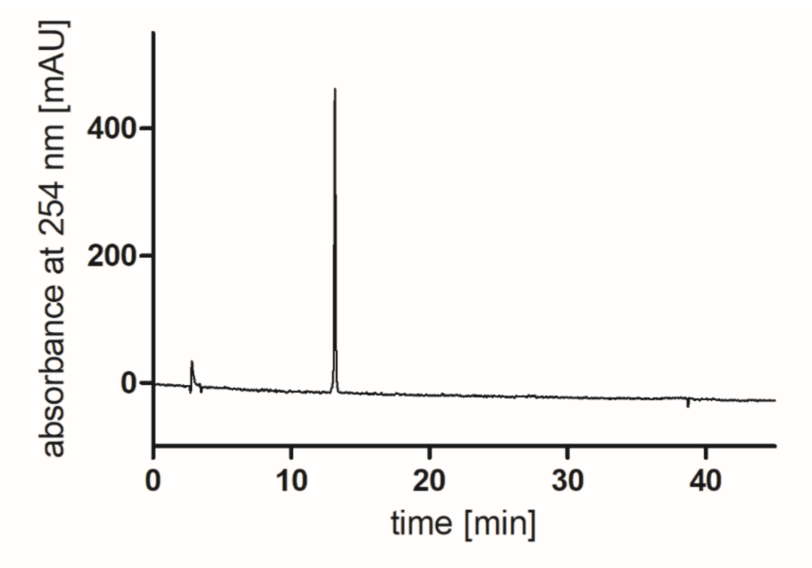


**Figure S3.** RP-HPLC analysis (purity control) of compound **20** (>99%, 254nm).

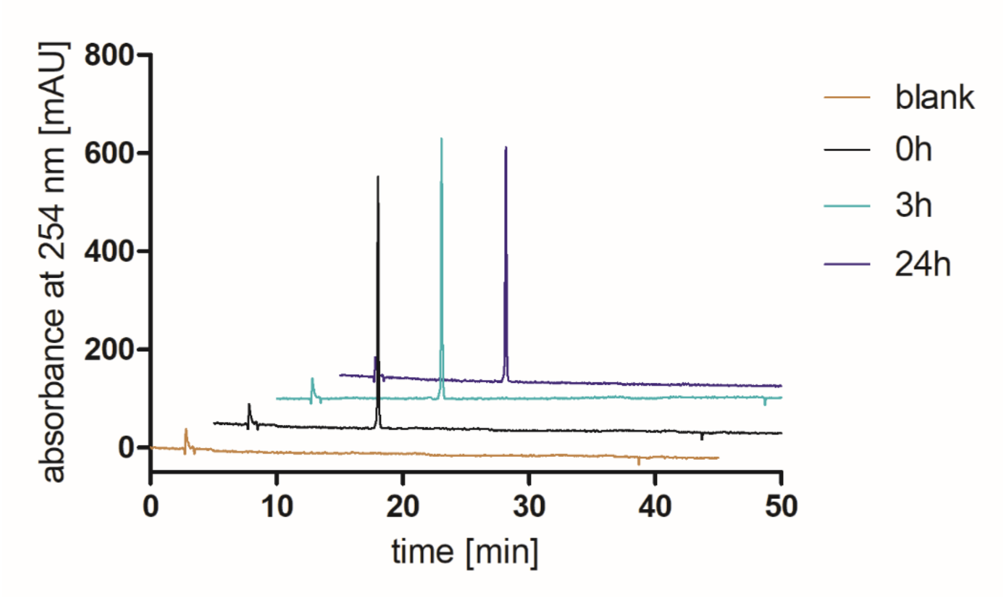


**Figure S4.** RP-HPLC analysis (stability control) of **20** after incubation in water/DMSO 1:1 at rt for up to 24 h. Compound **20** showed no decomposition.

### 2. Dopamine-induced $G_{o1}$ activation

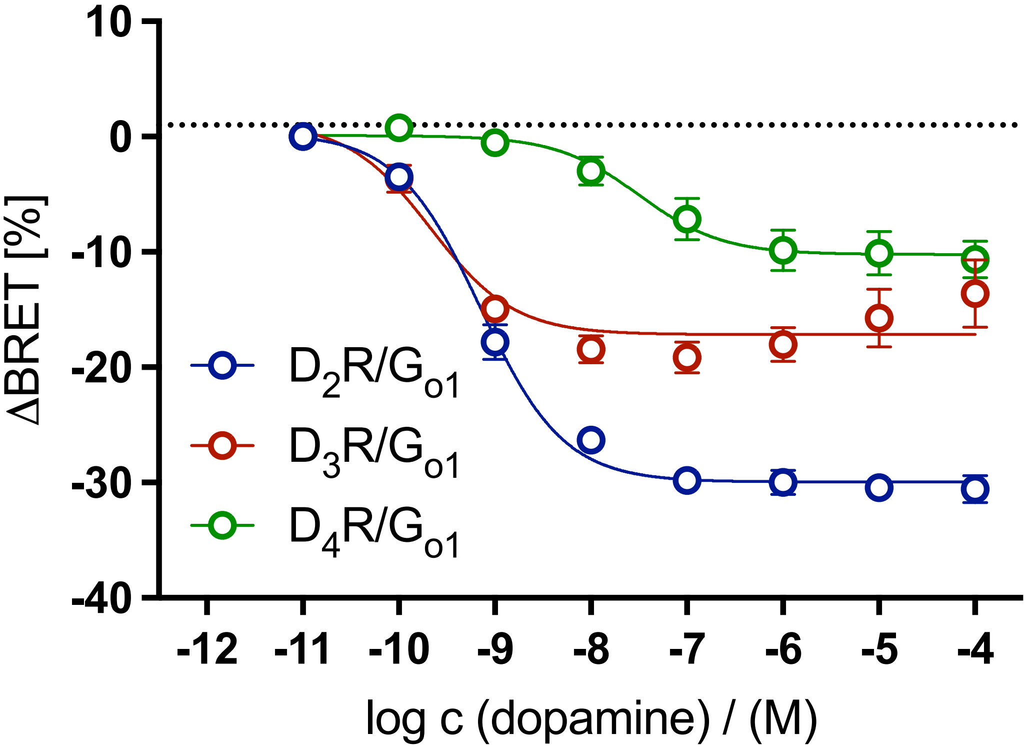


**Figure S5.** Concentration−response curves (CRCs) for $G_{o1}$ activation of dopamine in HEK293A cells transiently expressing the $G_{o1}$ BRET sensor along with the wild-type $D_2R$, $D_3R$ or $D_4R$. Graphs represent the means of three independent experiments each performed in duplicate. Data were analyzed by nonlinear regression and were best fitted to sigmoidal concentration-response curves.

### 3. NMR spectra

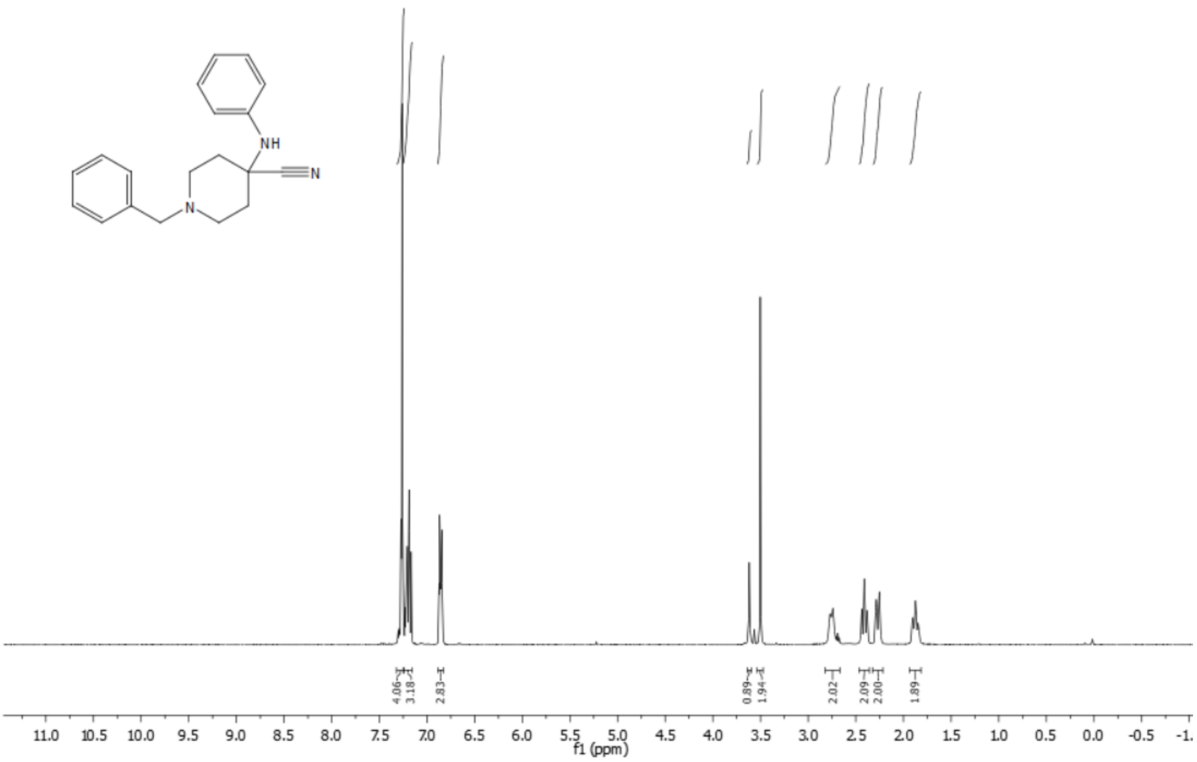

**Figure S6.** $^{1}H$ NMR spectrum (400 MHz, $CDCl_3$) of compound **2**.

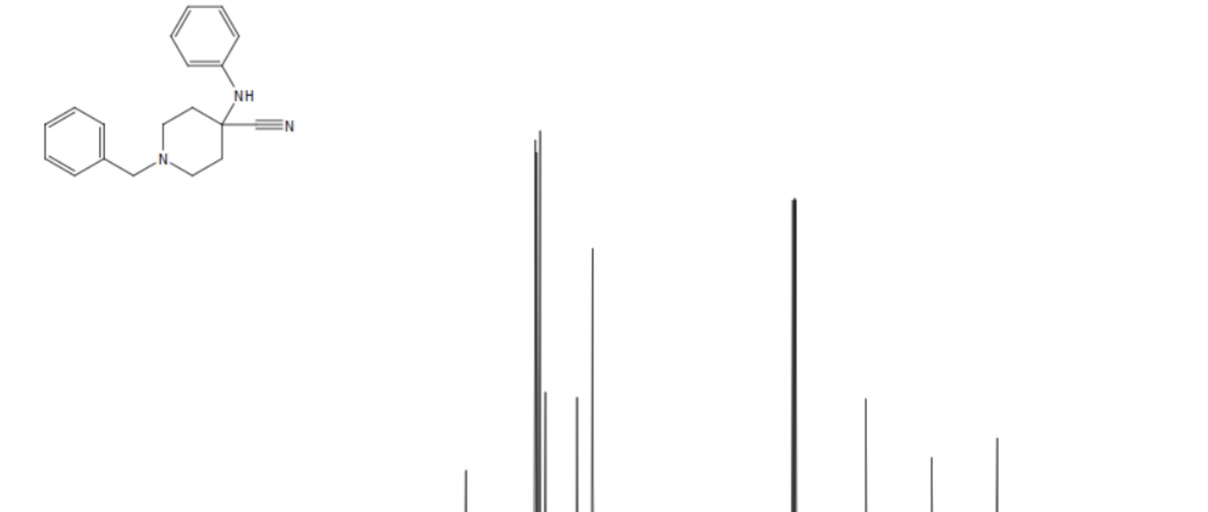

**Figure S7.** $^{13}C$ NMR spectrum (101 MHz, $CDCl_3$) of compound **2**.

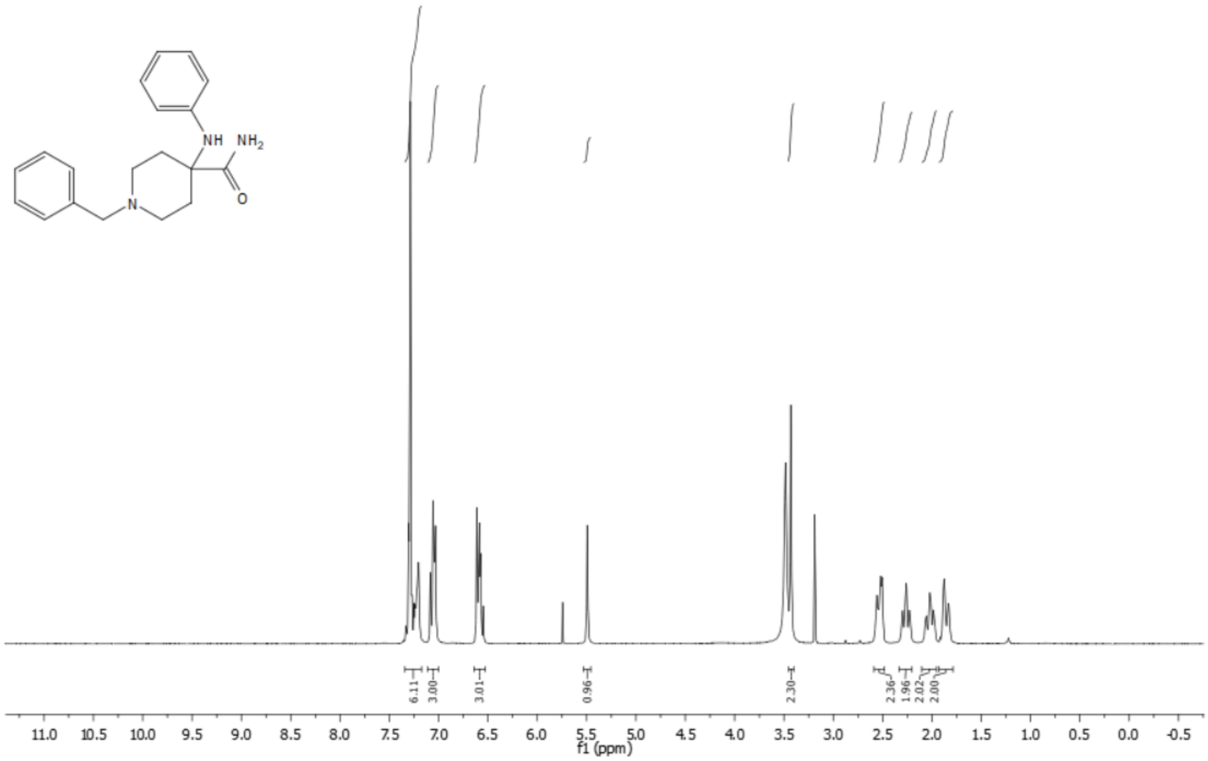


**Figure S8.** $^1$H NMR spectrum (300 MHz, DMSO-d$_6$) of compound **3**.

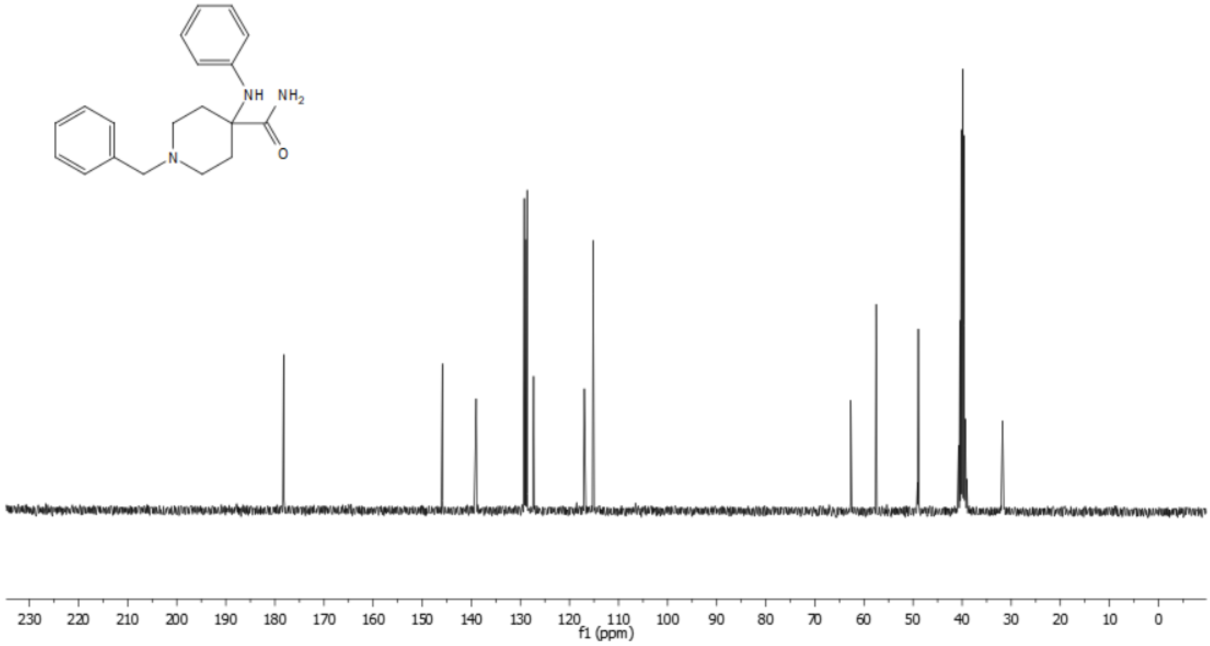


**Figure S9.** $^{13}$C NMR spectrum (101 MHz, DMSO-d$_6$) of compound **3**.

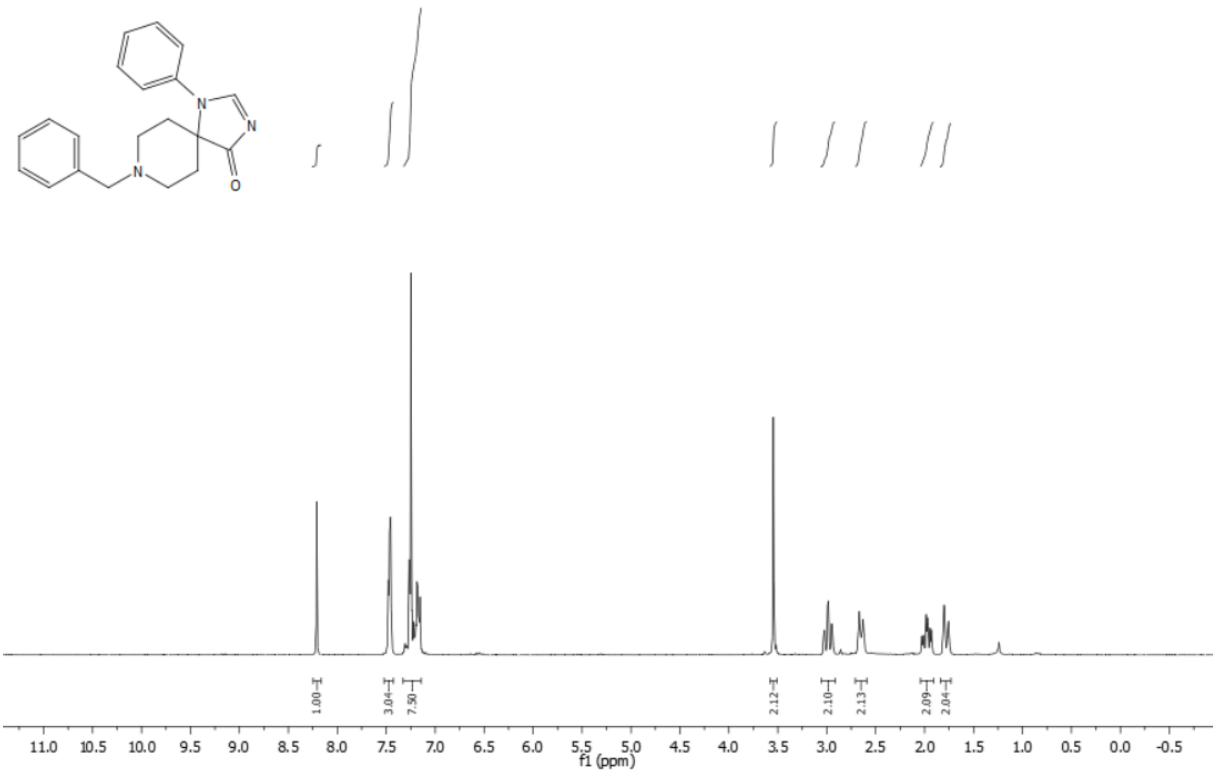

**Figure S10.** $^{1}$H NMR spectrum (300 MHz, $CDCl_3$) of compound **4**.

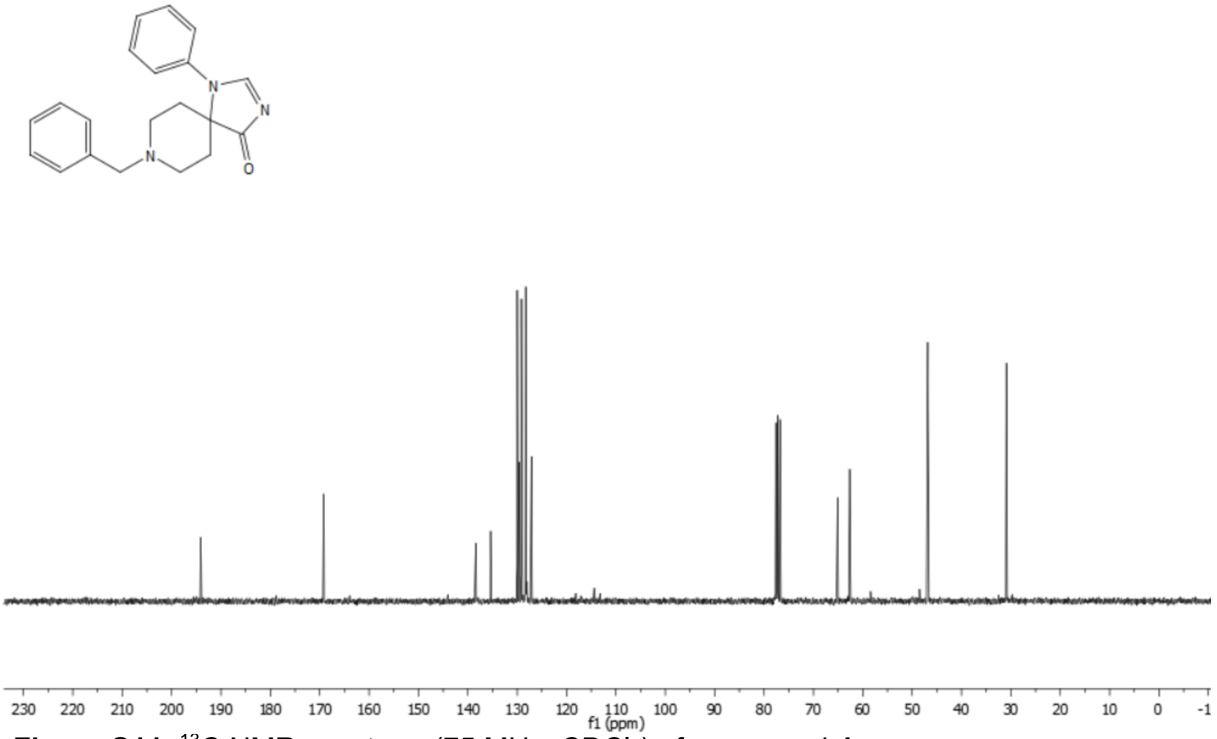

**Figure S11.** $^{13}$C NMR spectrum (75 MHz, $CDCl_3$) of compound **4**.

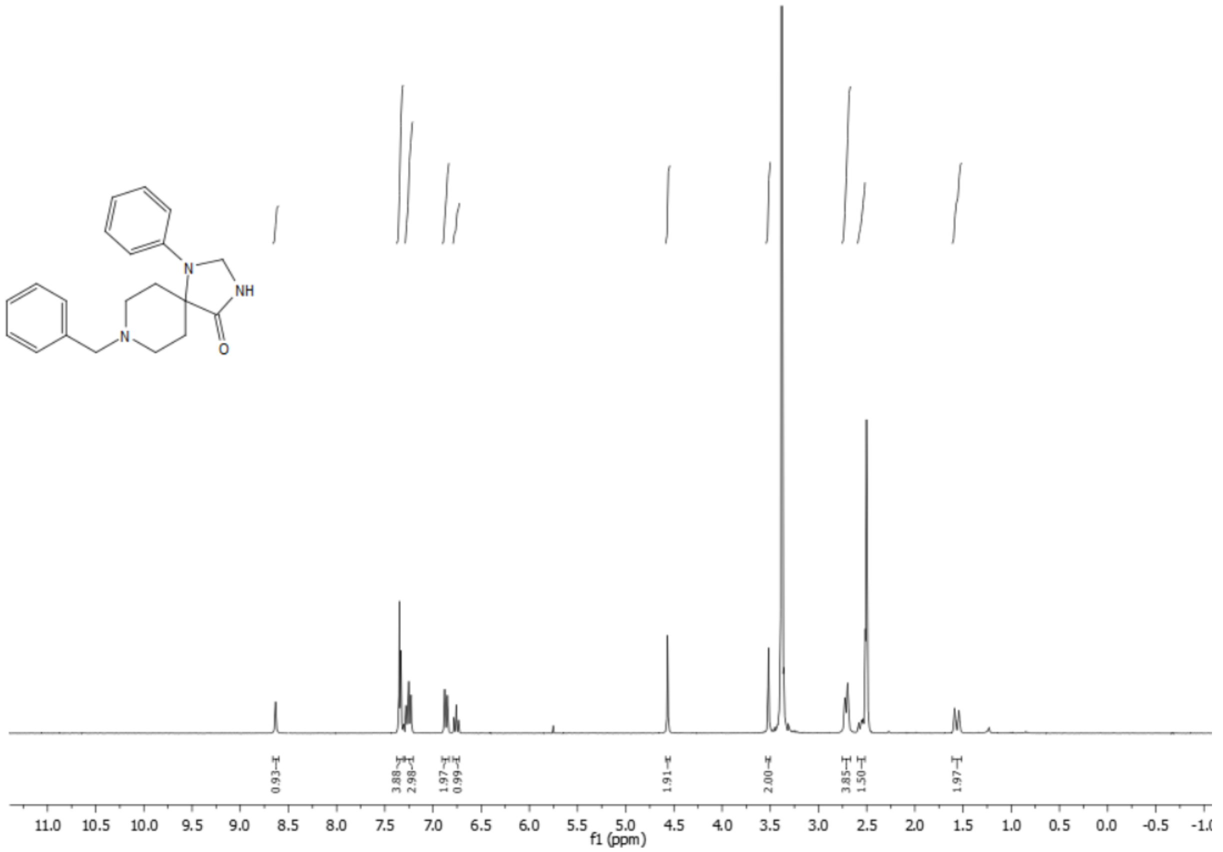


**Figure S12.** $^{1}H$ NMR spectrum (300 MHz, DMSO-$d_6$) of compound **5**.

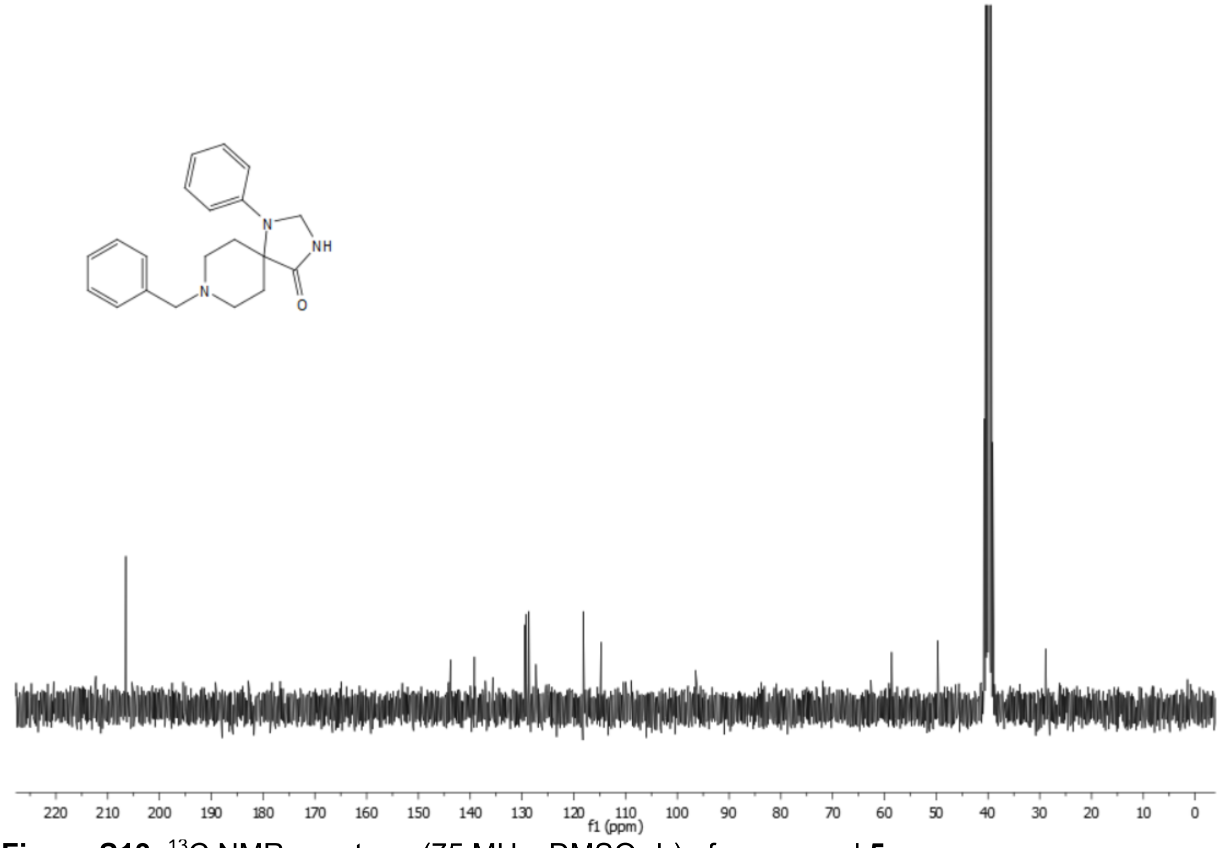


**Figure S13.** $^{13}C$ NMR spectrum (75 MHz, DMSO-$d_6$) of compound **5**.

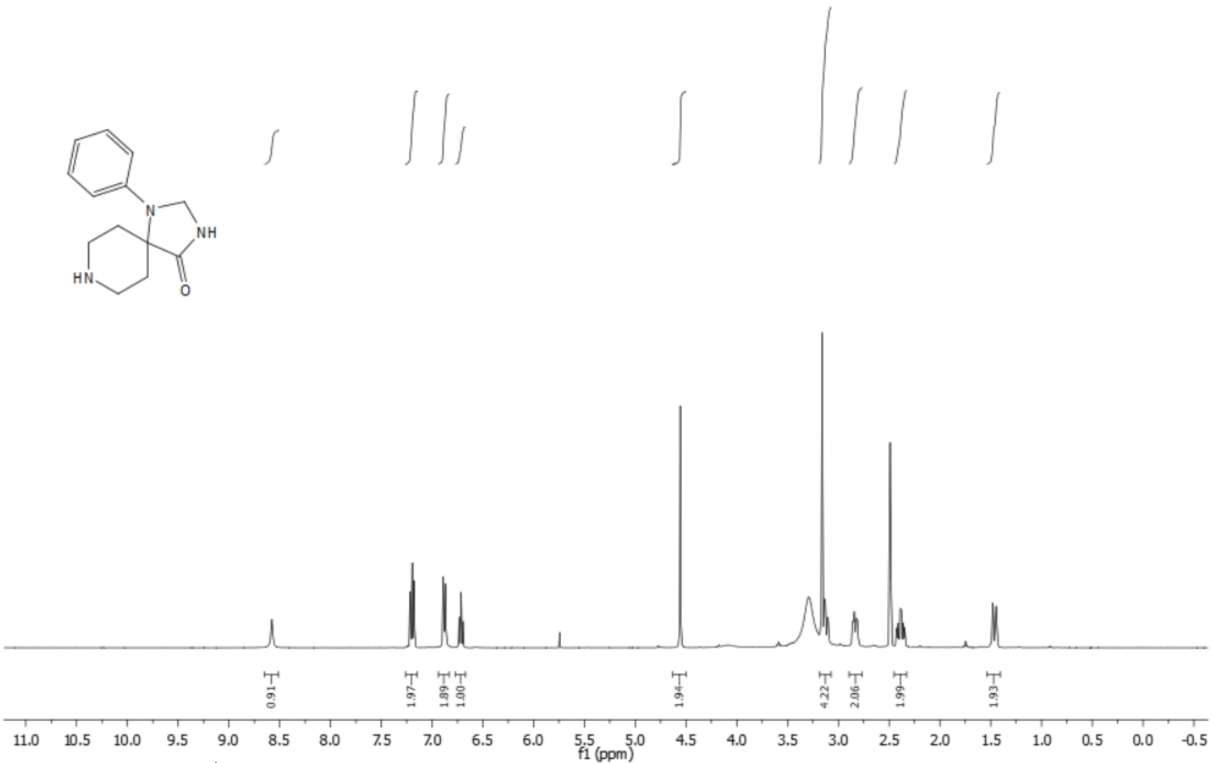


**Figure S14.** $^{1}$H NMR spectrum (400 MHz, DMSO-$d_6$) of compound **6**.

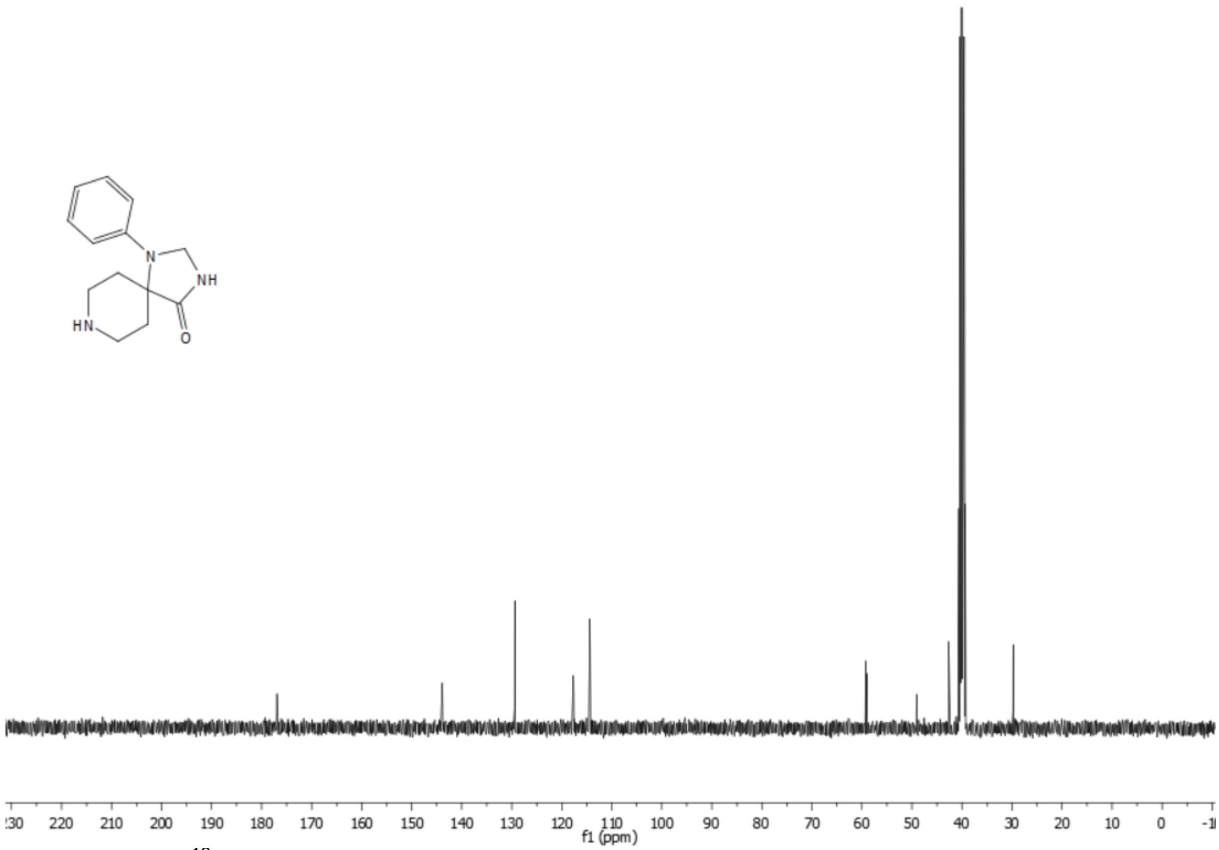


**Figure S15.** $^{13}$C NMR spectrum (101 MHz, DMSO-$d_6$) of compound **6**.

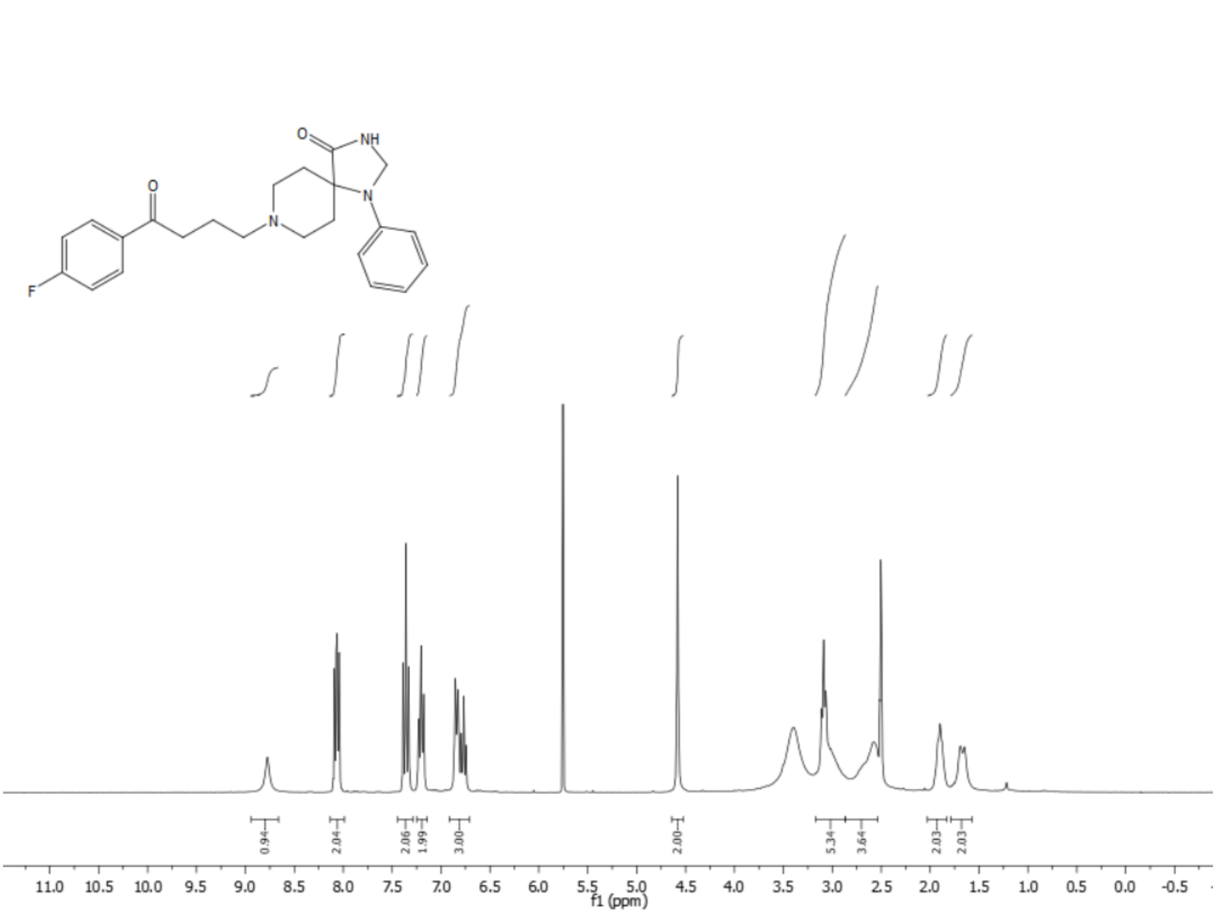


**Figure S16.** $^{1}$H NMR spectrum (400 MHz, DMSO-$d_6$) of compound **7**.

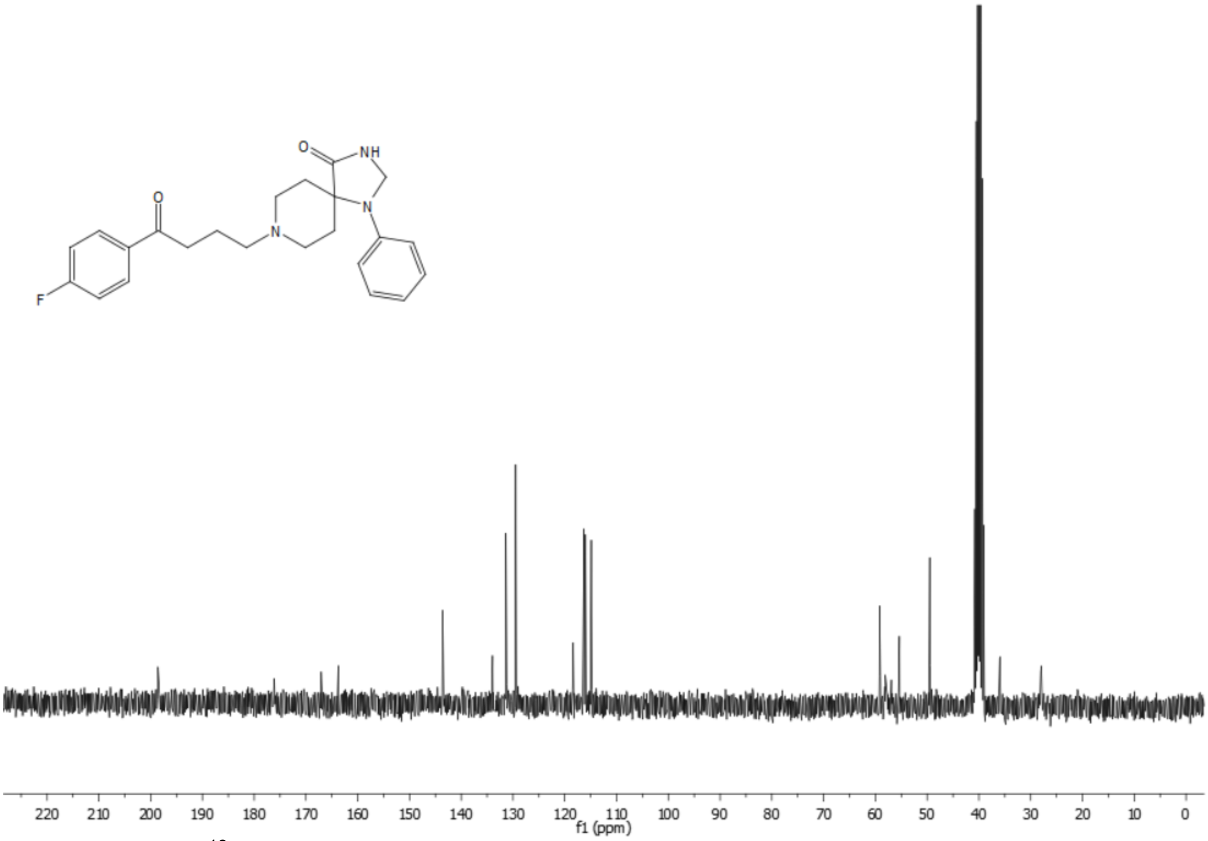


**Figure S17.** $^{13}$C NMR spectrum (101 MHz, DMSO-$d_6$) of compound **7**.

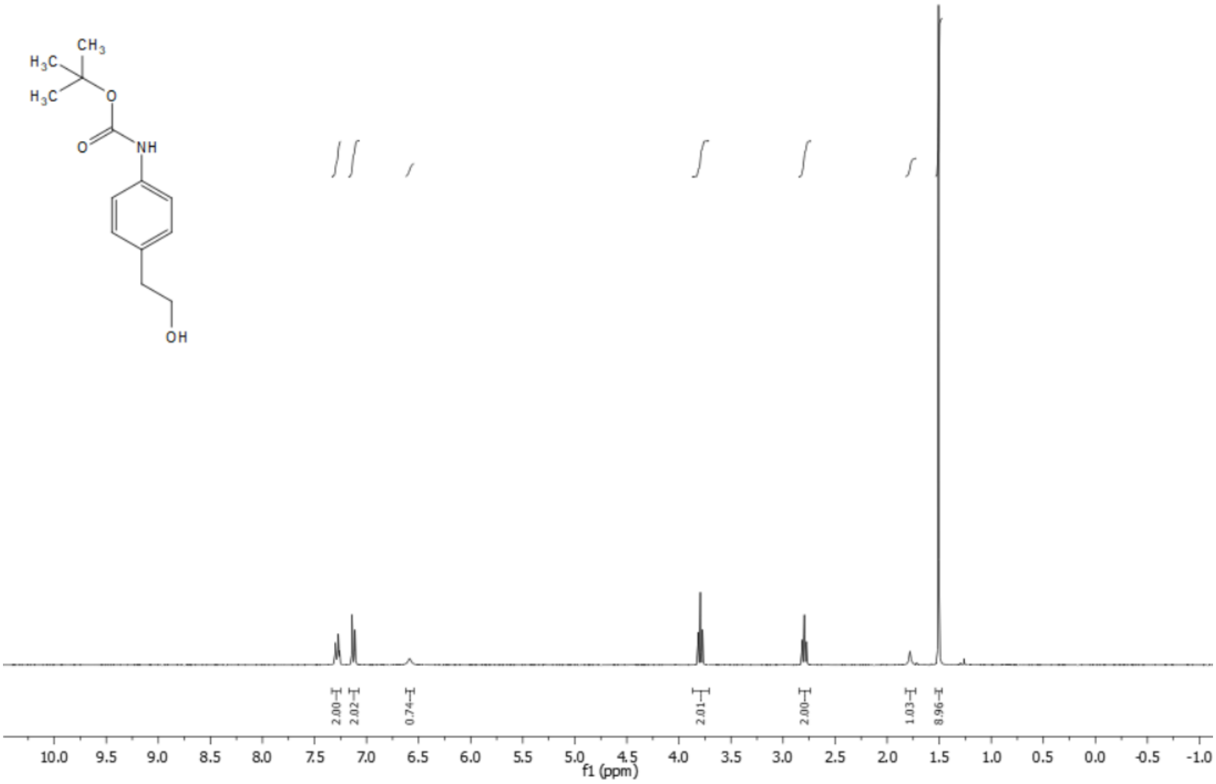


**Figure S18.** $^{1}H$ NMR spectrum (300 MHz, $CDCl_3$) of compound **8**.

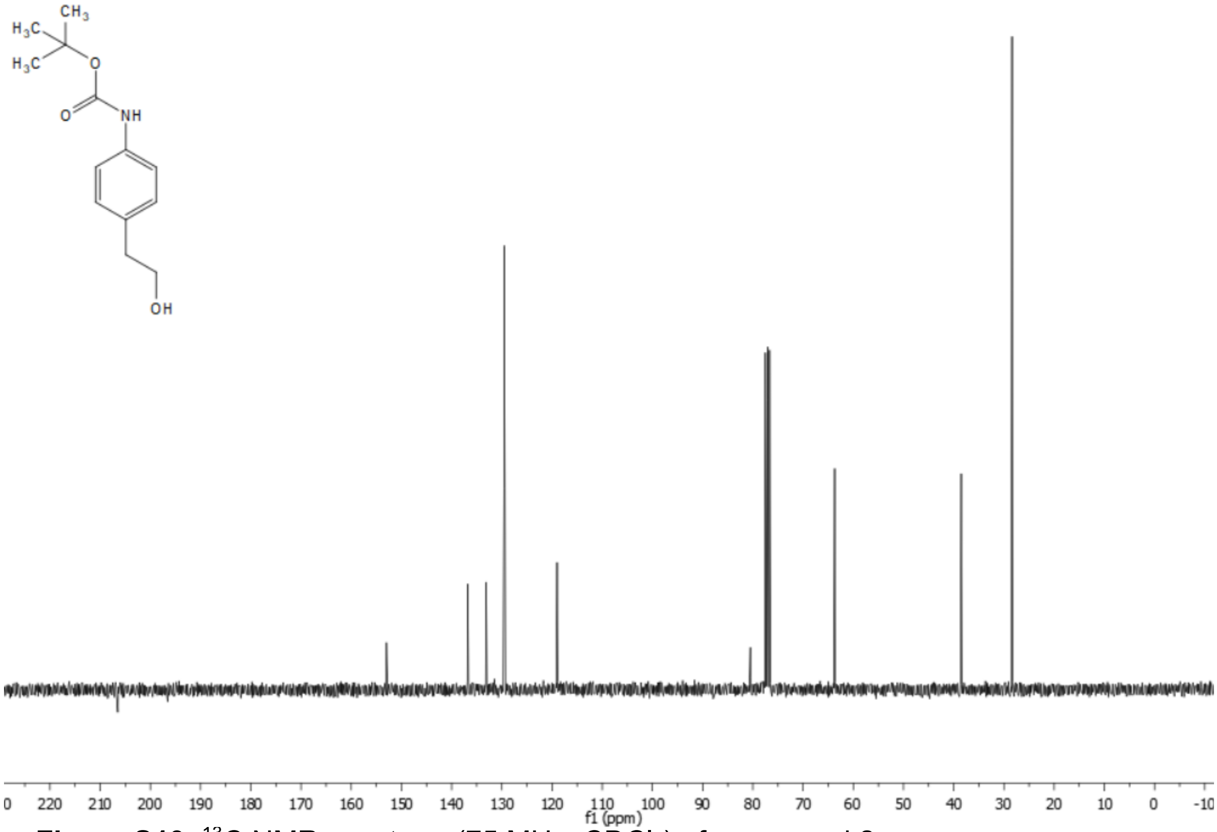


**Figure S19.** $^{13}C$ NMR spectrum (75 MHz, $CDCl_3$) of compound **8**.

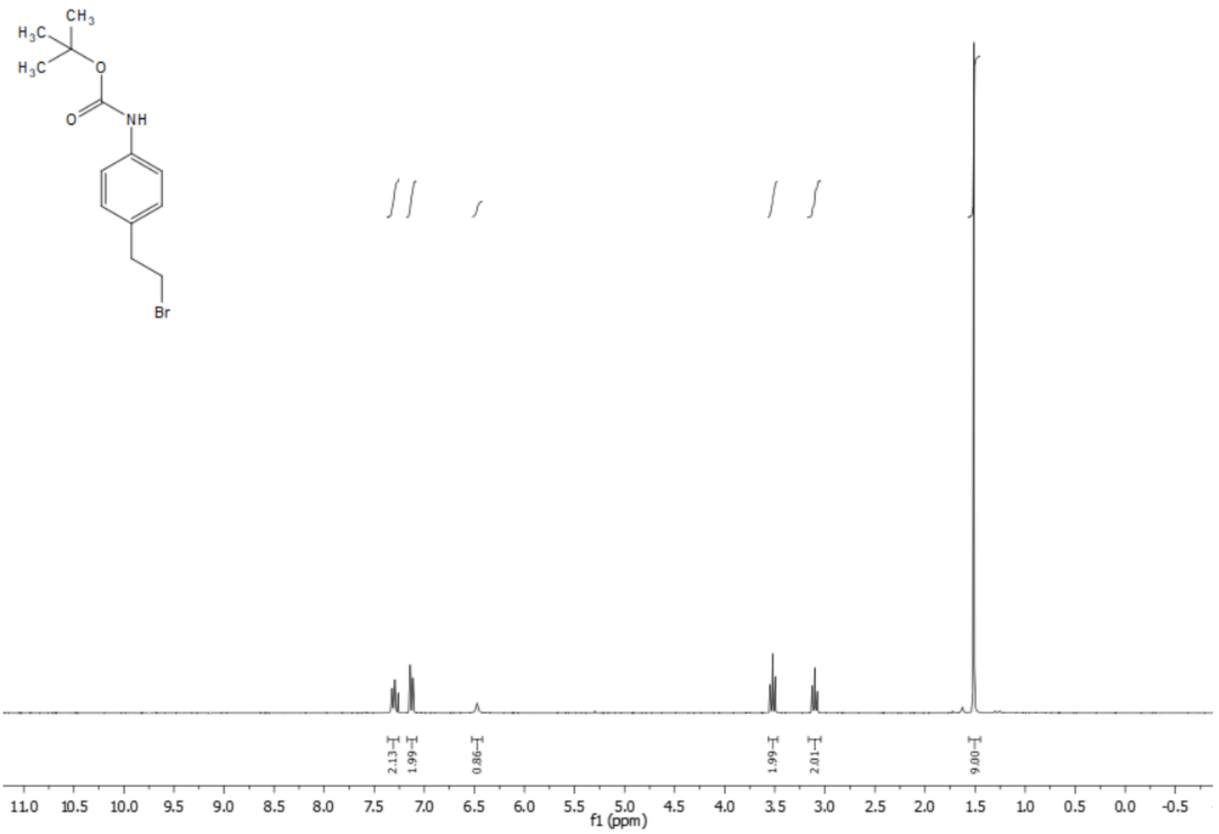


**Figure S20.** $^{1}H$ NMR spectrum (300 MHz, $CDCl_3$) of compound **9**.

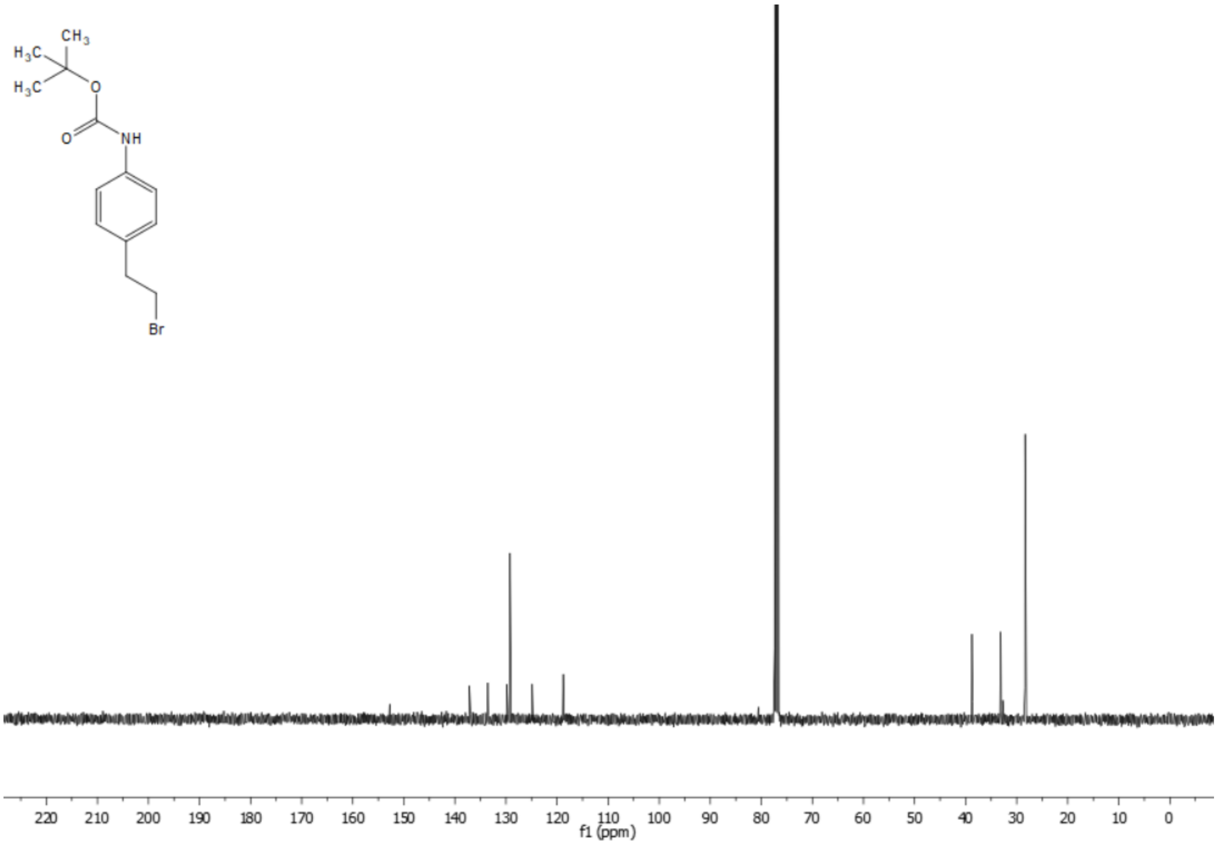


**Figure S21.** $^{13}C$ NMR spectrum (75 MHz, $CDCl_3$) of compound **9**.

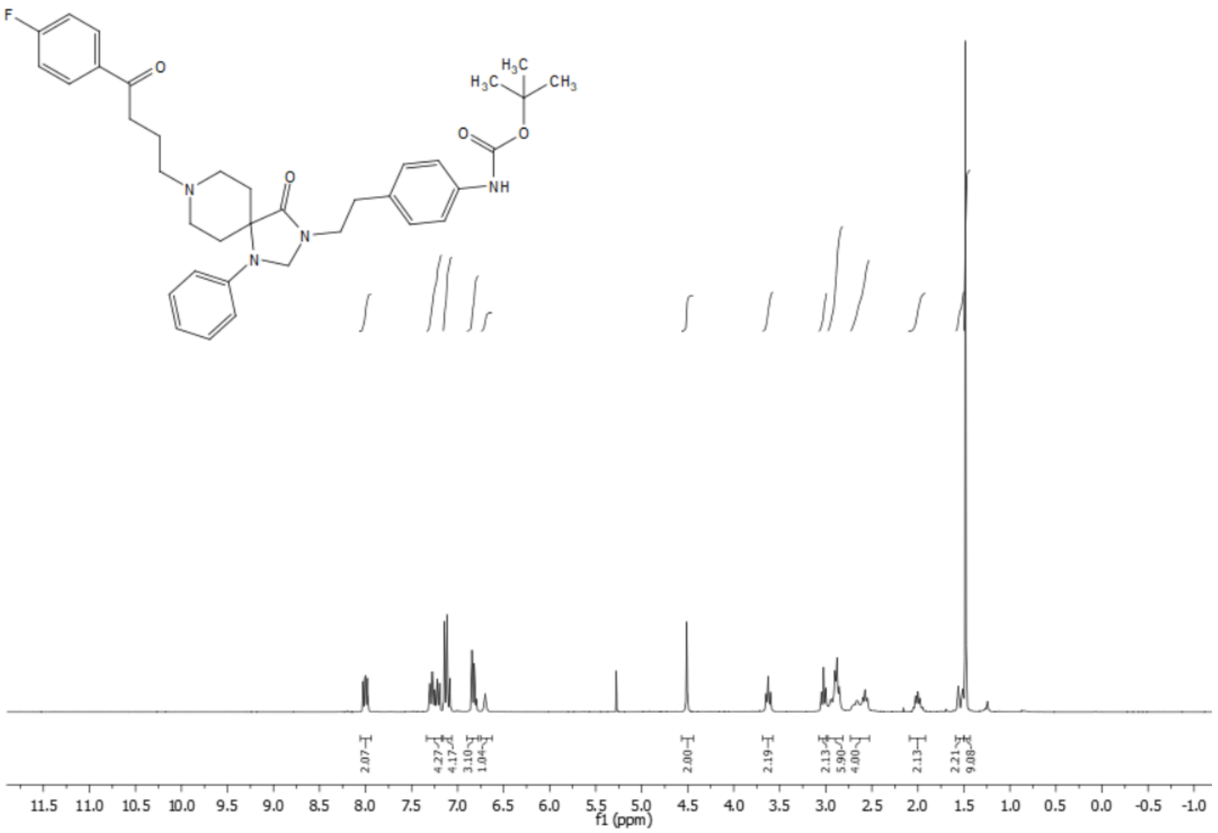


**Figure S22.** $^{1}$H NMR spectrum (400 MHz, $CDCl_3$) of compound **10**.

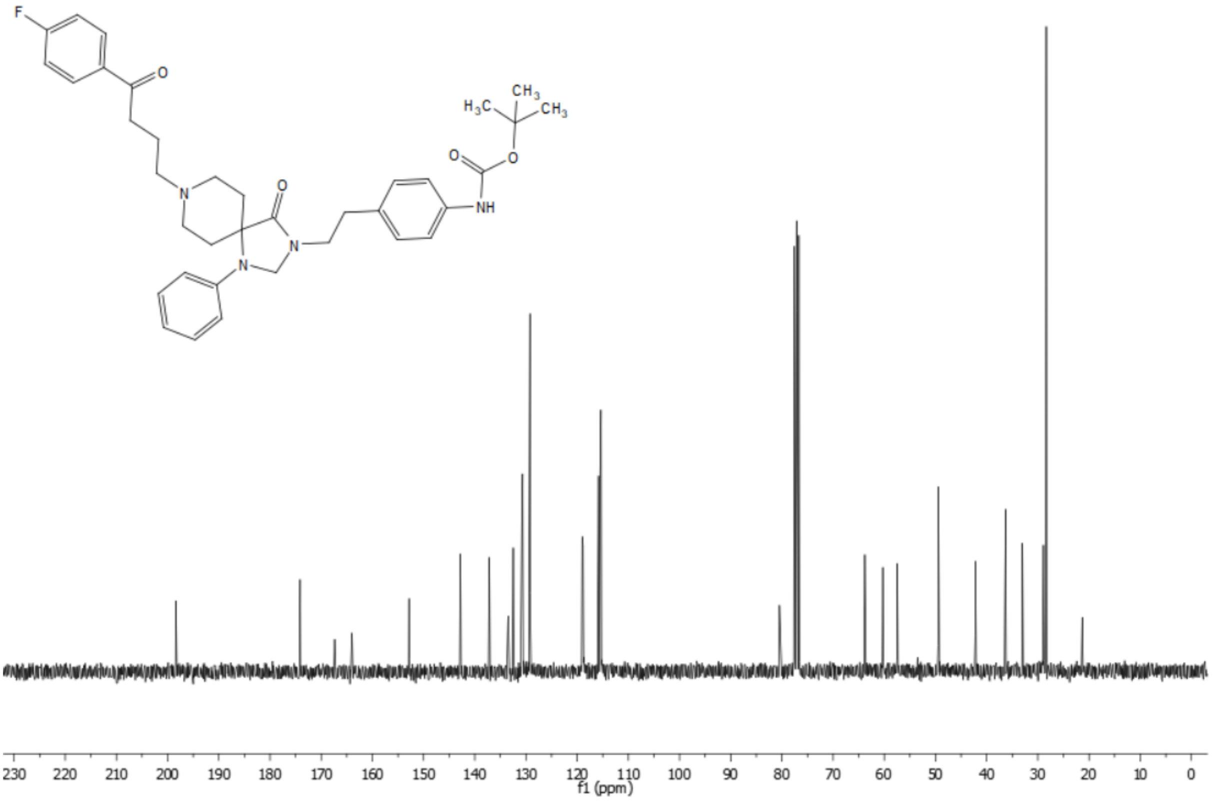


**Figure S23.** $^{13}$C NMR spectrum (75 MHz $CDCl_3$) of compound **10**.

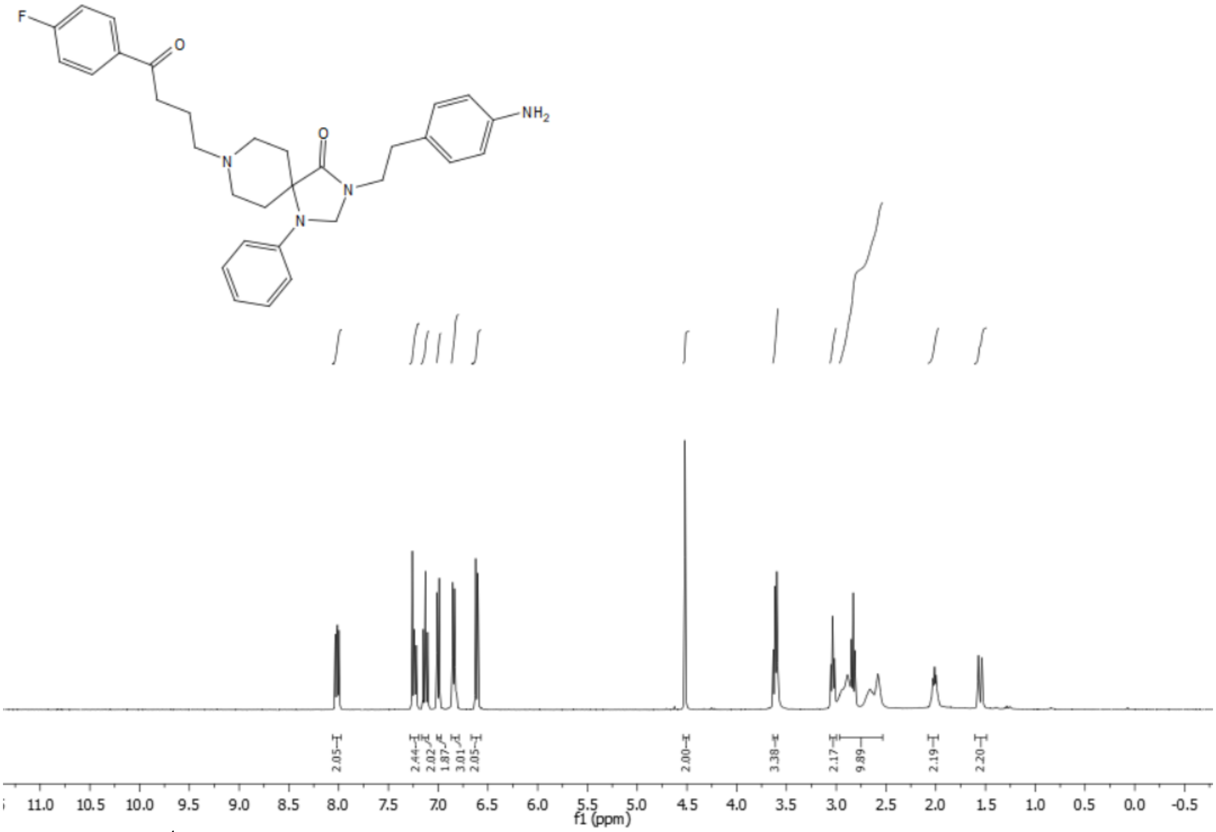


**Figure S24.** $^{1}H$ NMR spectrum (400 MHz, $CDCl_3$) of compound **11**.

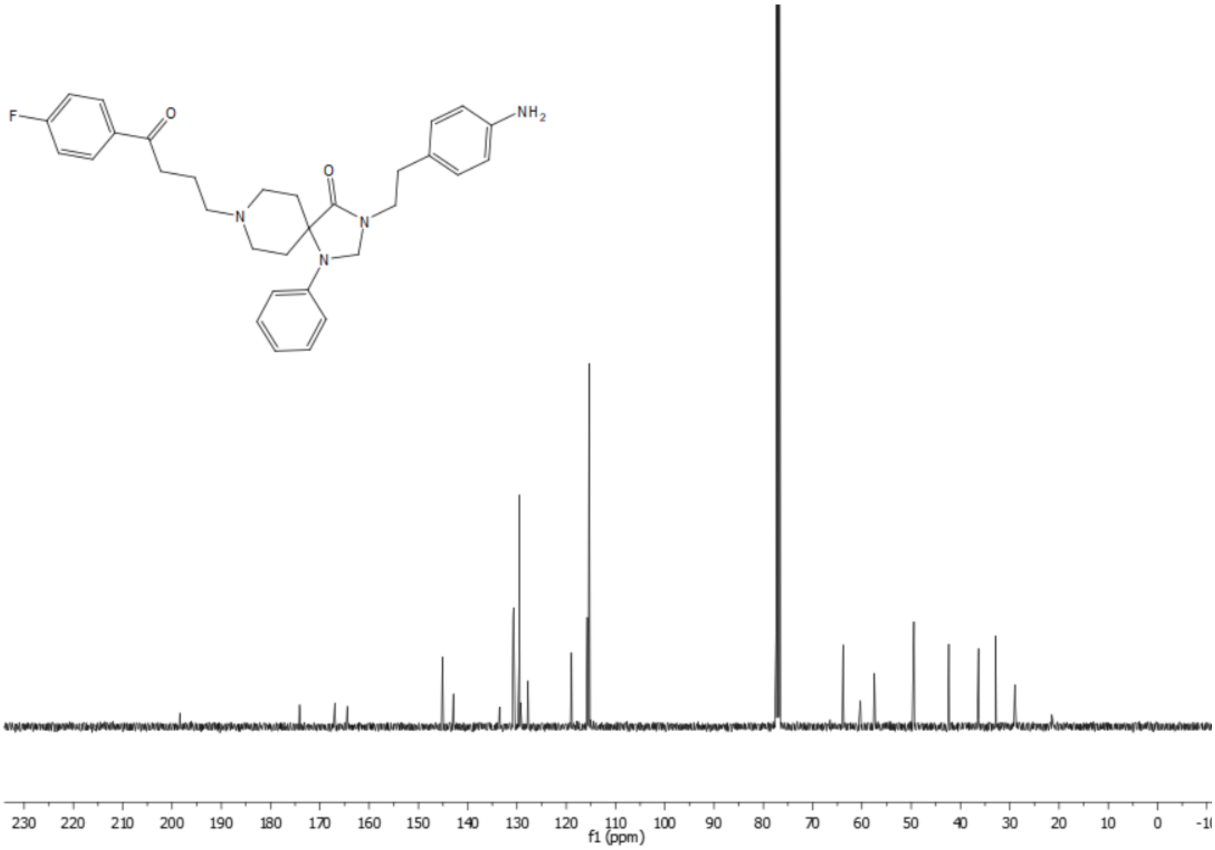


**Figure S25.** $^{13}C$ NMR spectrum (101 MHz, $CDCl_3$) of compound **11**.

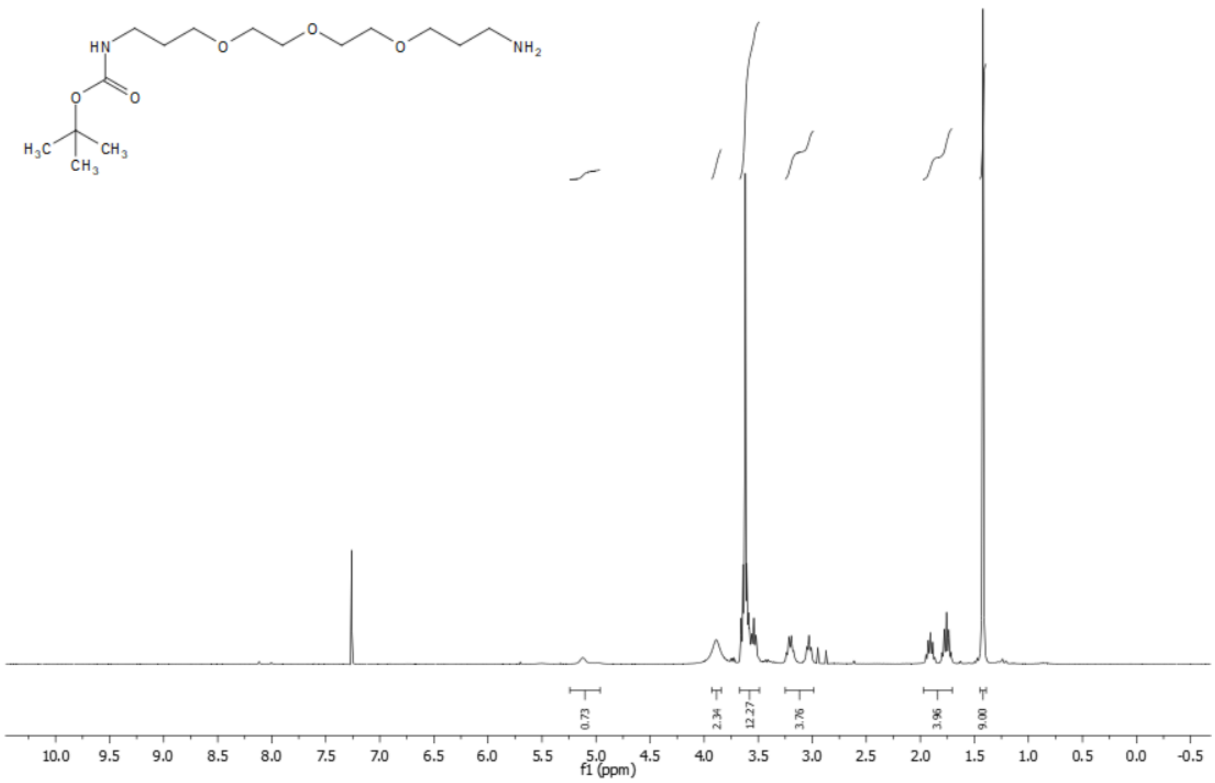

**Figure S26.** $^{1}$H NMR spectrum (300 MHz, $CDCl_3$) of compound **12**.

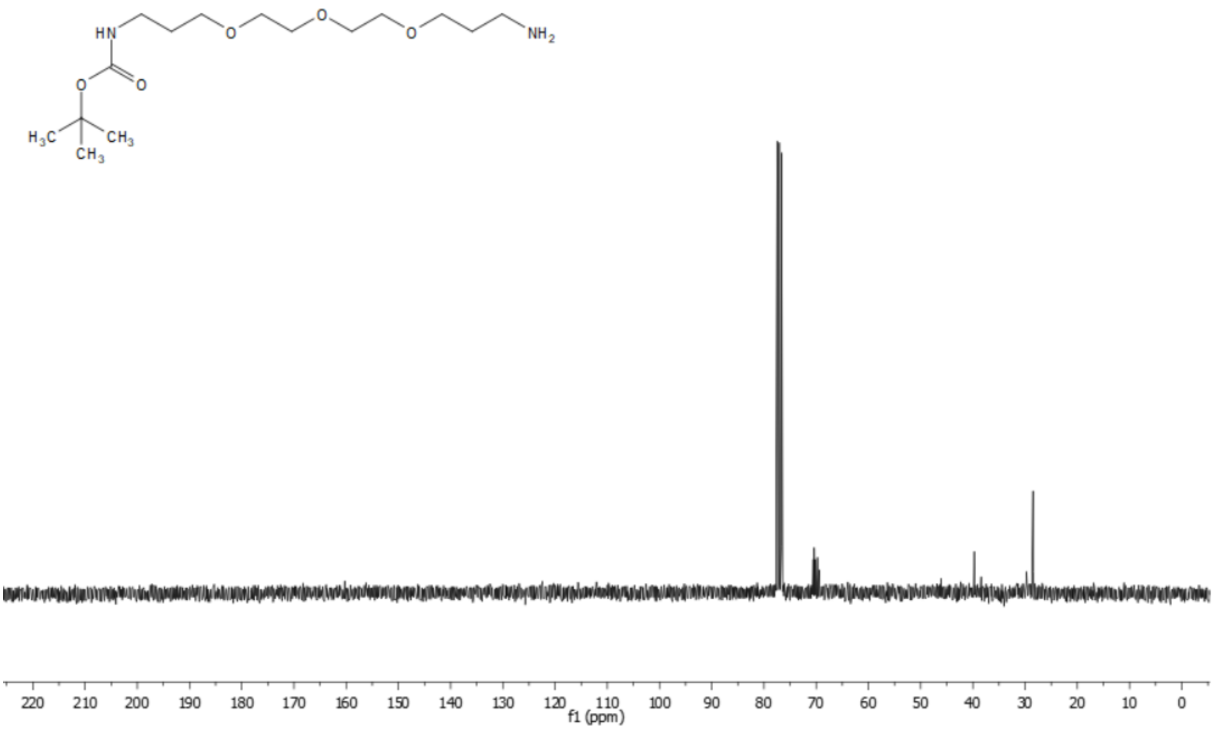

**Figure S27.** $^{13}$C NMR spectrum (75 MHz, $CDCl_3$) of compound **12**.

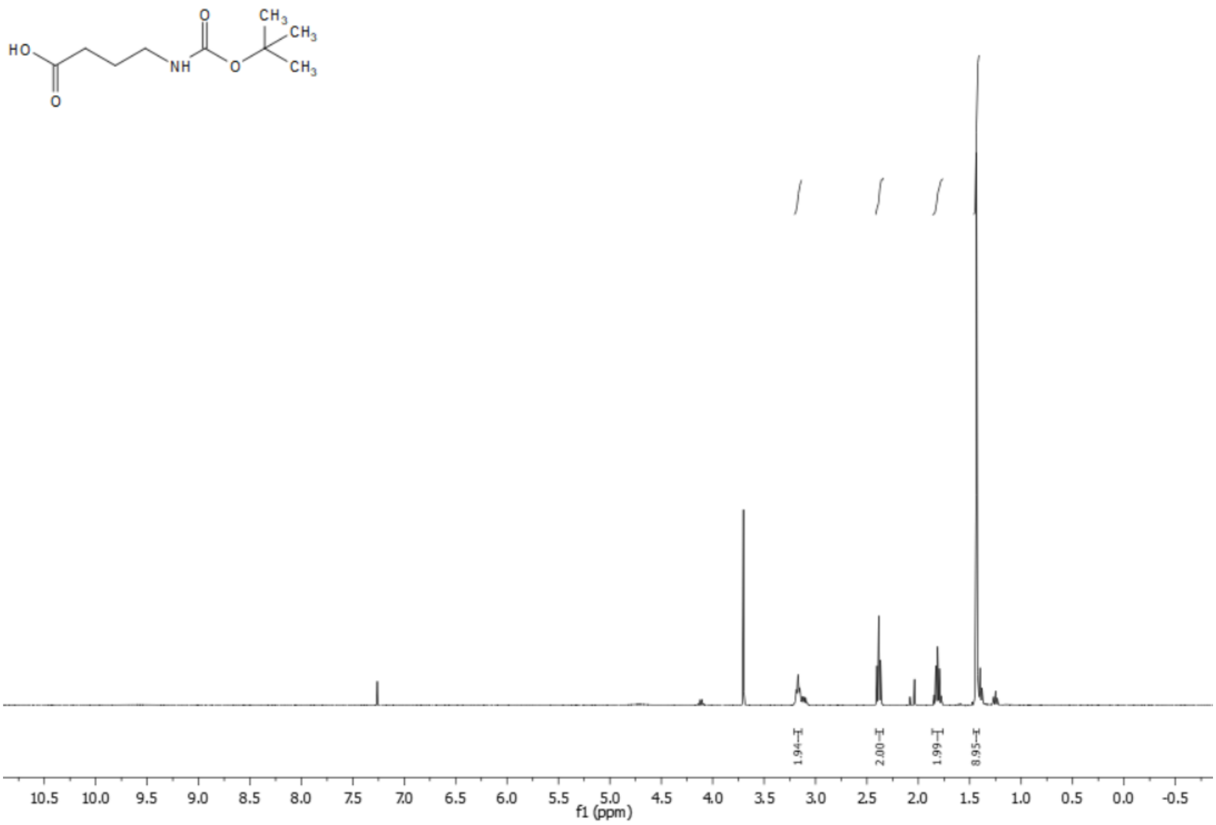


**Figure S28.** $^{1}$H NMR spectrum (400 MHz, $CDCl_3$) of compound **13**.

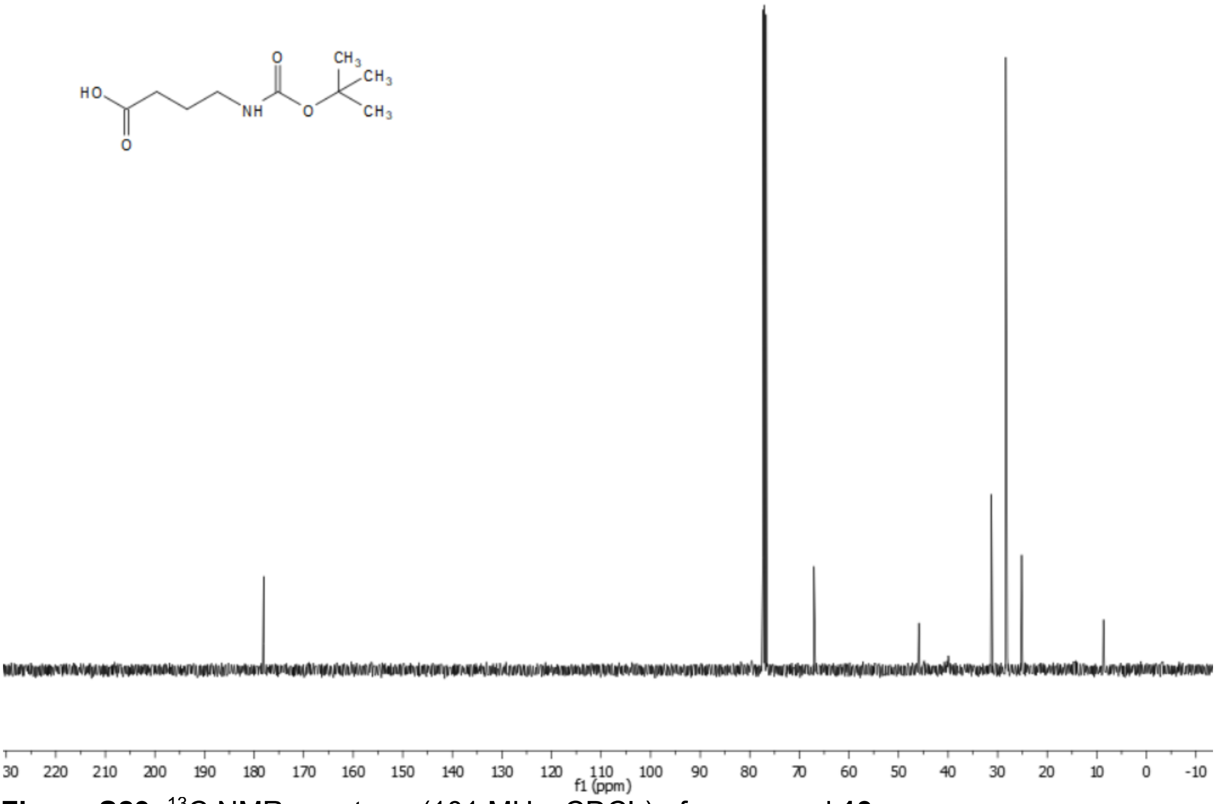


**Figure S29.** $^{13}$C NMR spectrum (101 MHz, $CDCl_3$) of compound **13**.

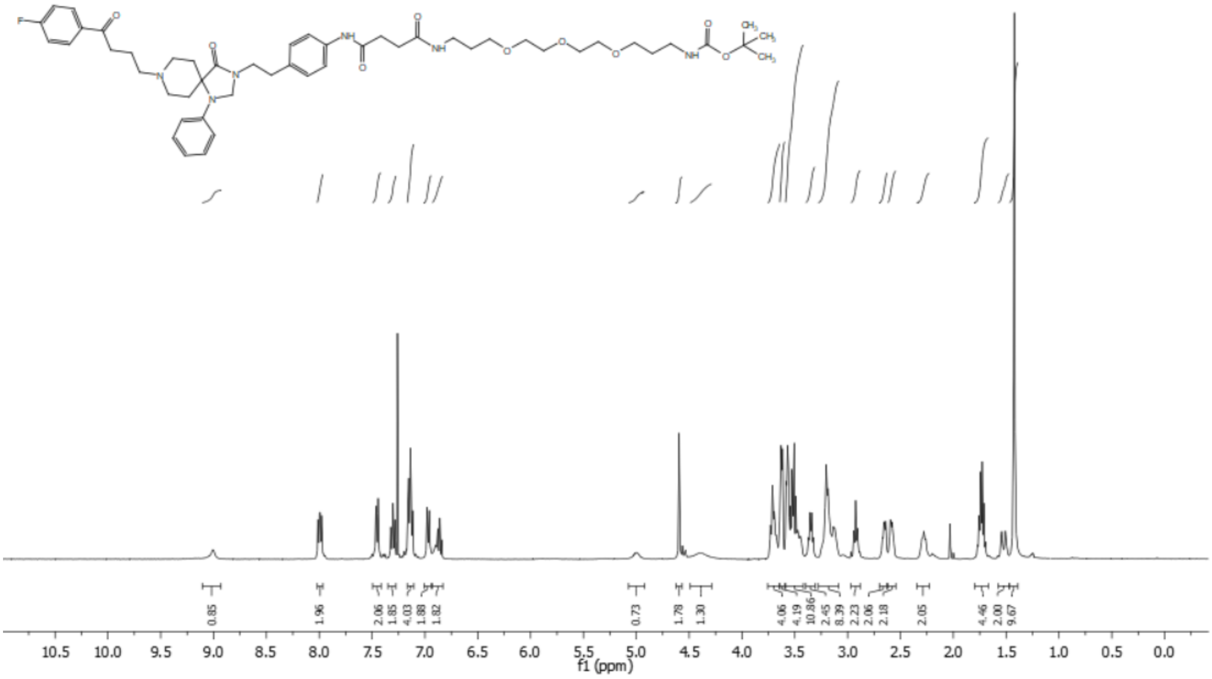


**Figure S30.** $^{1}H$ NMR spectrum (400 MHz, $CDCl_3$) of compound **14**.

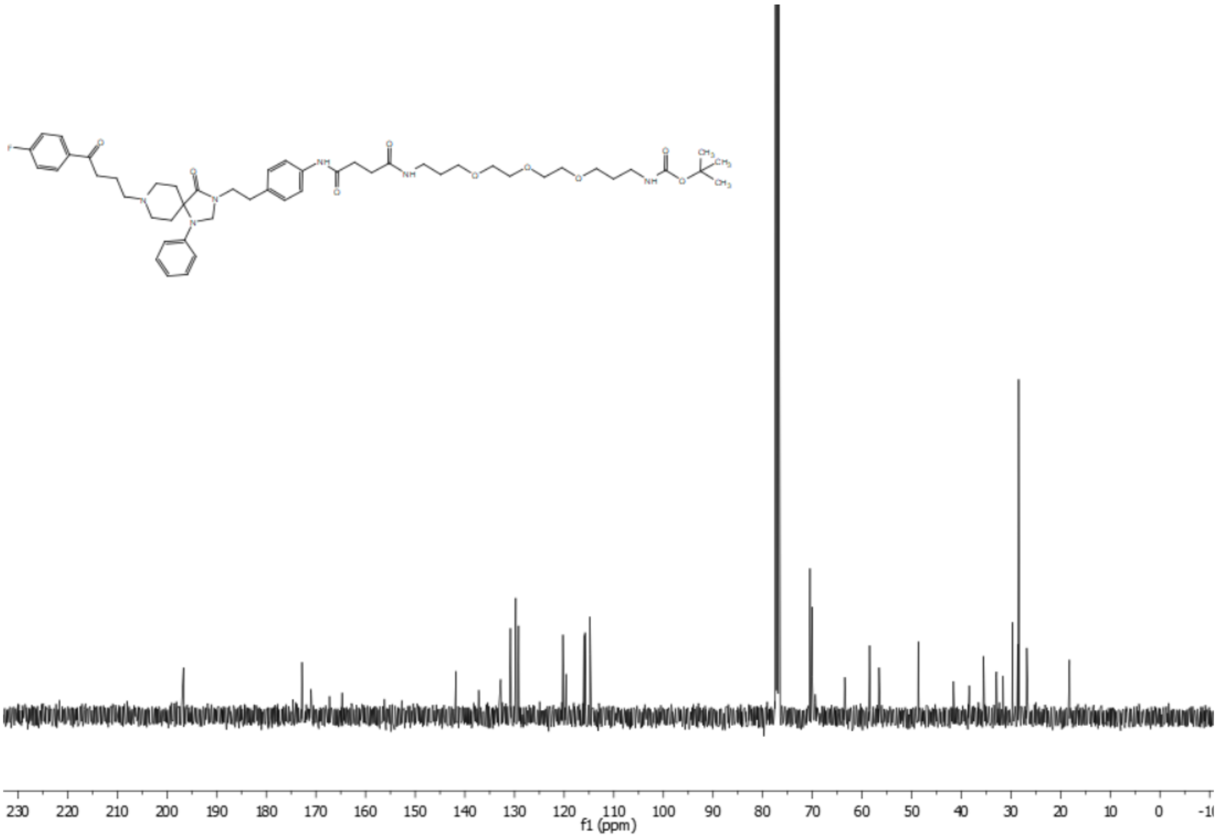


**Figure S31.** $^{13}C$ NMR spectrum (400 MHz, $CDCl_3$) of compound **14**.

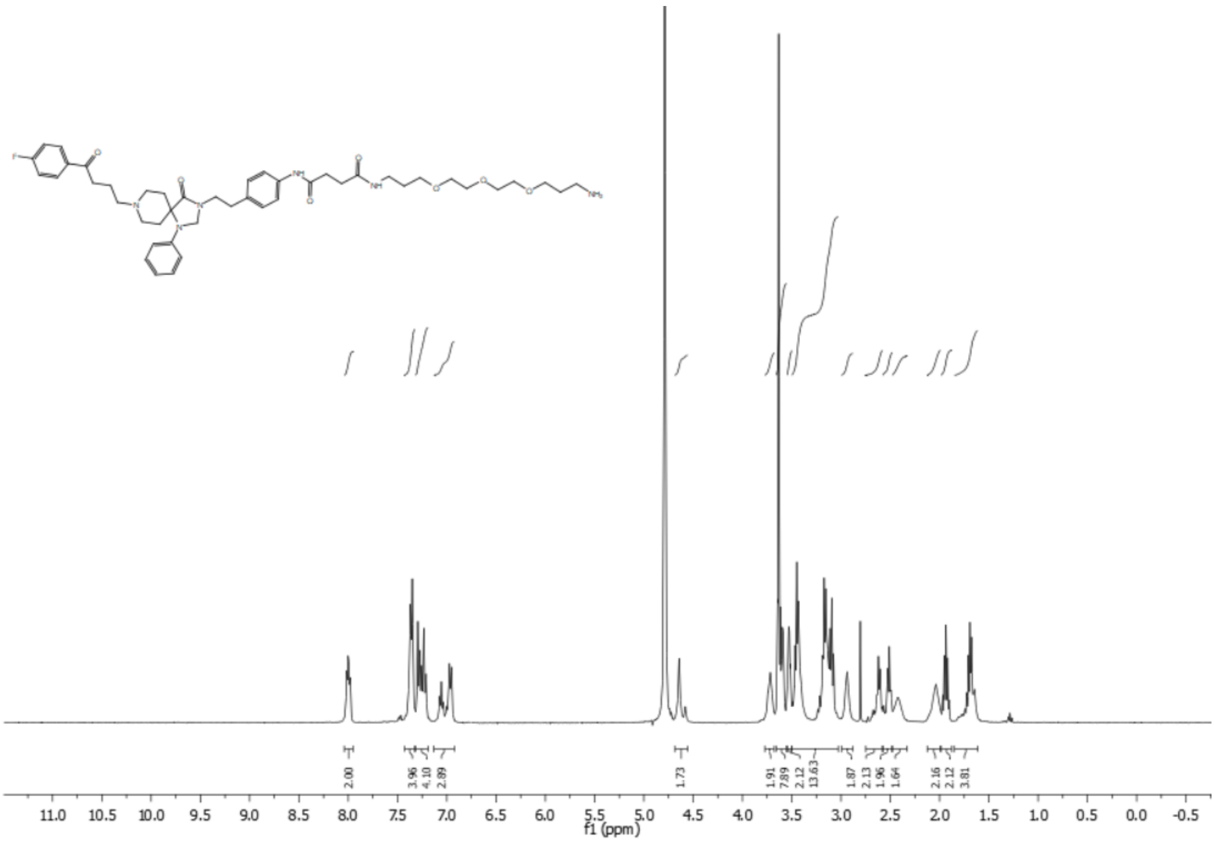


**Figure 32.** $^{1}H$ NMR spectrum (400 MHz, $D_2O$) of compound **15**.

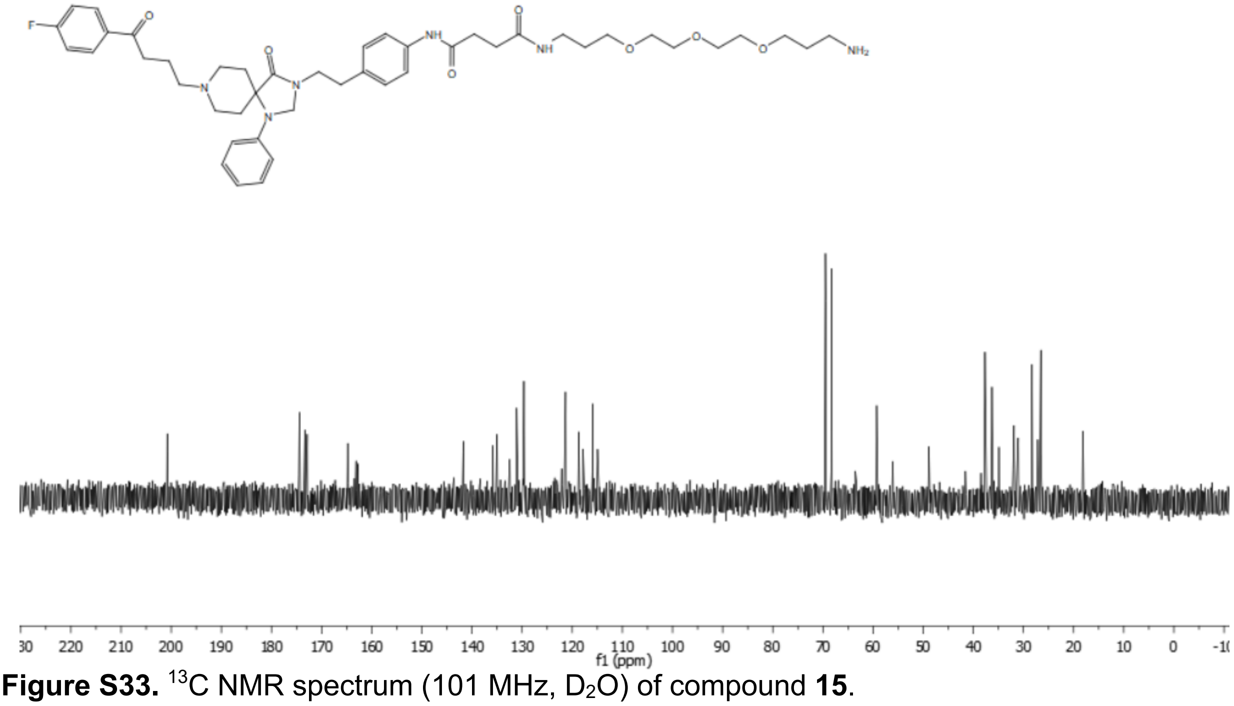


**Figure S33.** $^{13}C$ NMR spectrum (101 MHz, $D_2O$) of compound **15**.

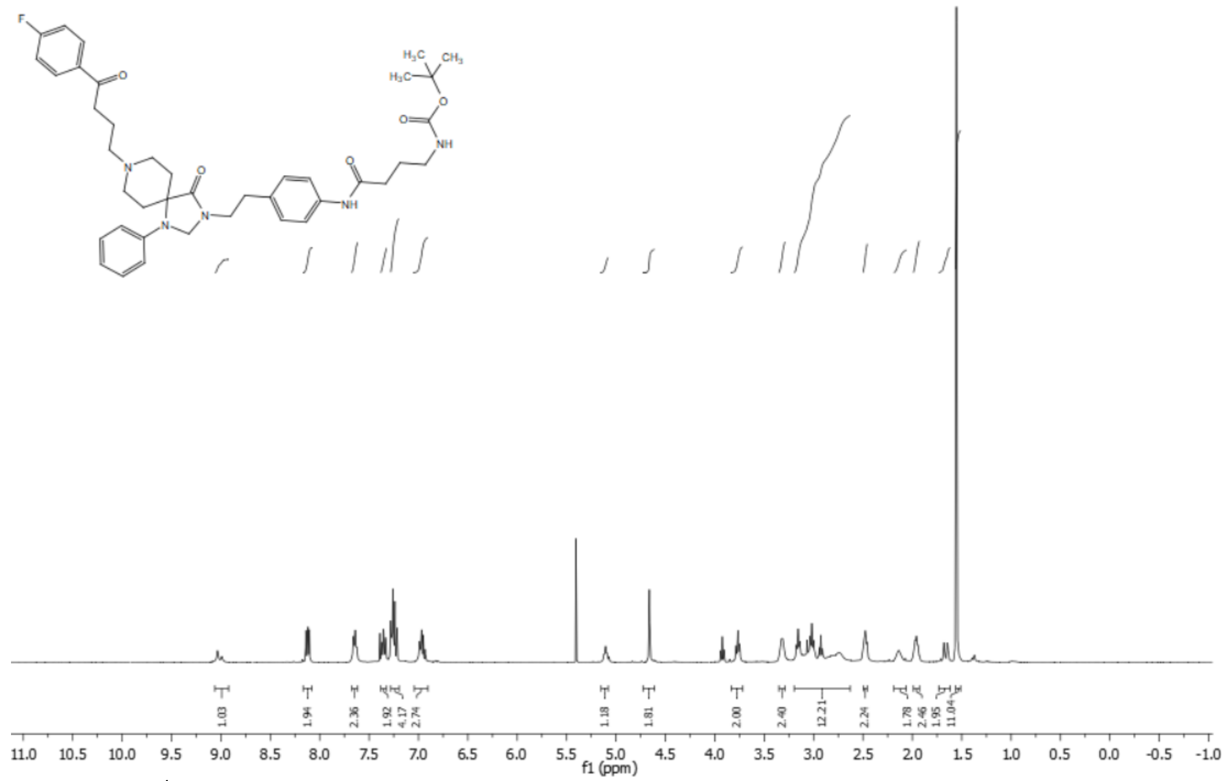

**Figure S34.** $^{1}$H NMR spectrum (400 MHz, $CDCl_3$) of compound **18**.

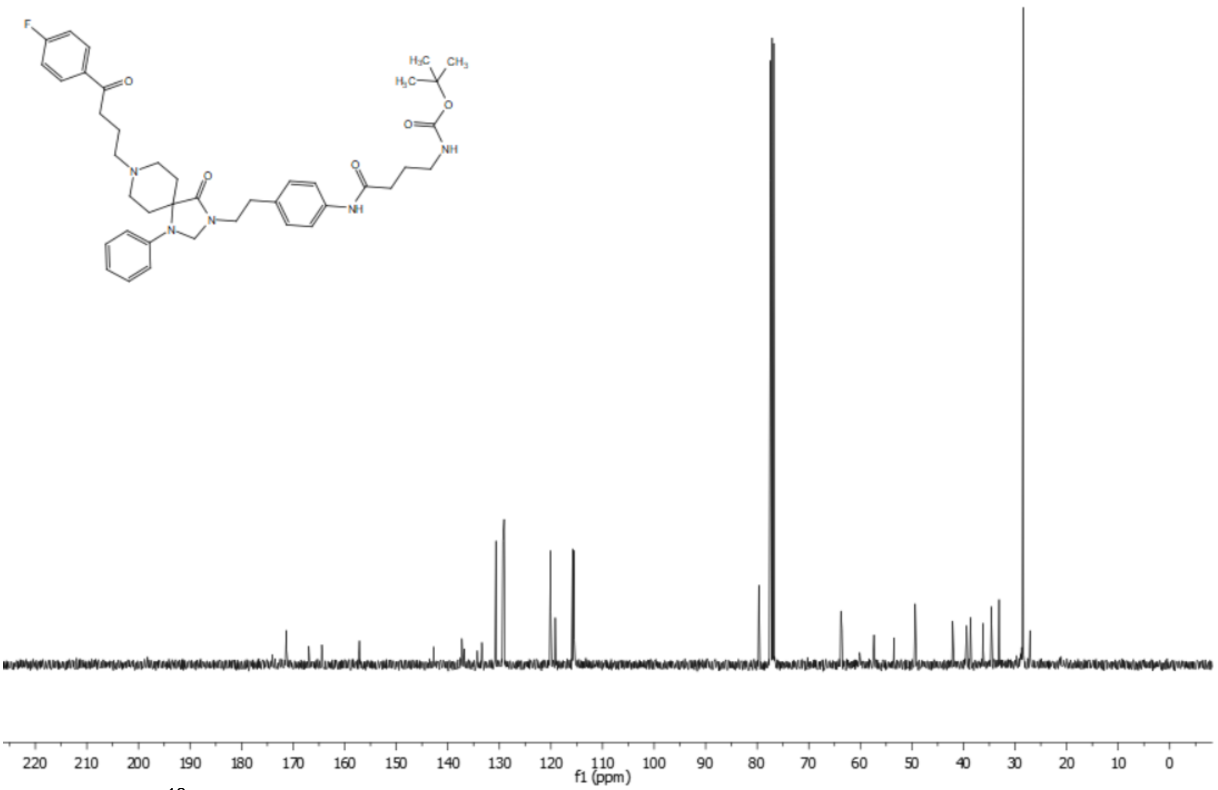

**Figure S35.** $^{13}$C NMR spectrum (101 MHz, $CDCl_3$) of compound **18**.

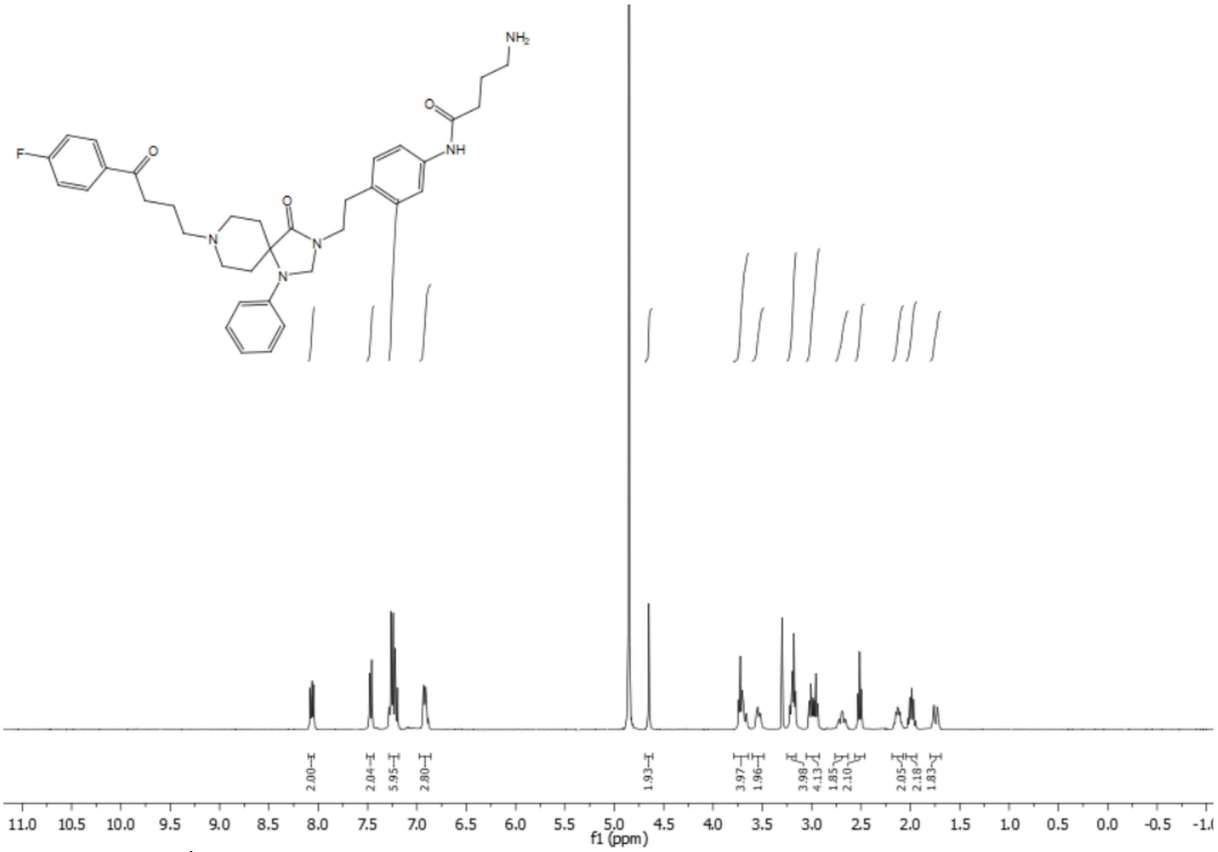

**Figure S36.** $^{1}$H NMR spectrum (400 MHz, $CD_3OD$) of compound **19**.

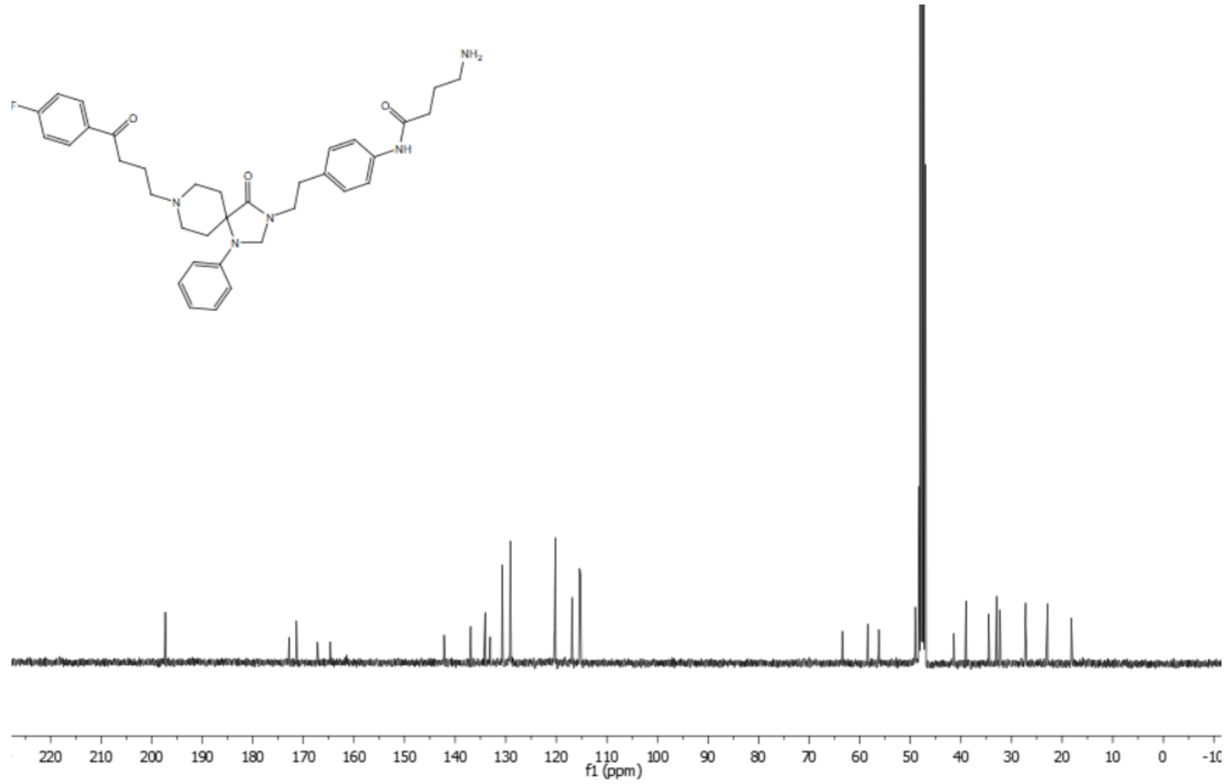

**Figure S37.** $^{13}$C NMR spectrum (101 MHz, $CD_3OD$) of compound **19**.

### 4. Structures of the fluorescent ligands 16, 17 and 20

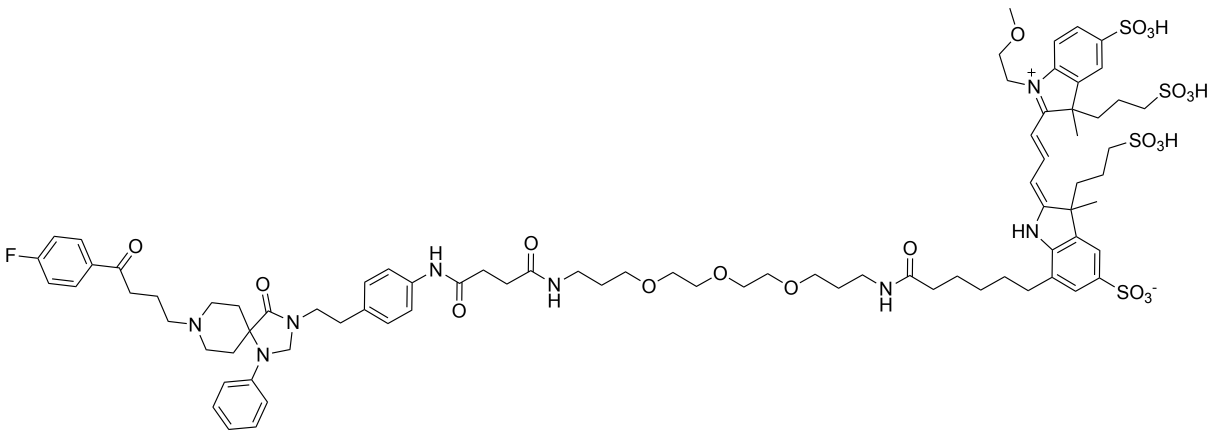


**Figure S38.** Structure of compound **16**.

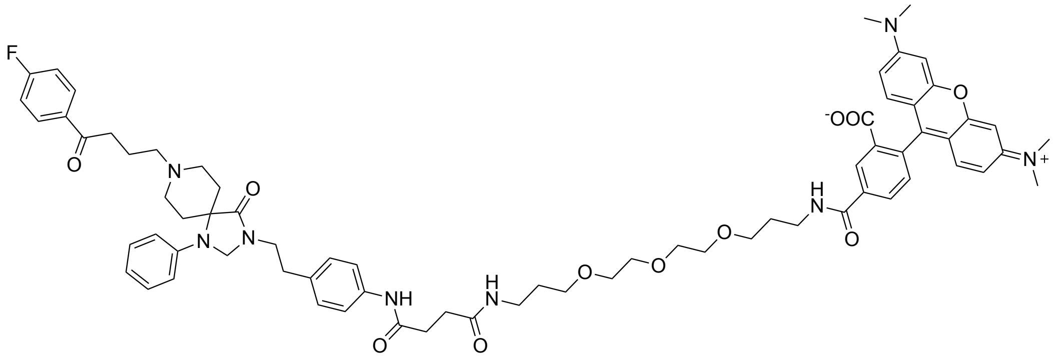


**Figure S39.** Structure of compound **17**.

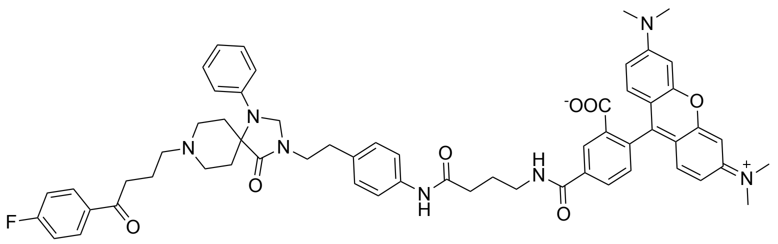


**Figure S40.** Structure of compound **20**.
